# Th17 cell master transcription factor RORC2 regulates HIV-1 gene expression and viral outgrowth

**DOI:** 10.1101/2021.03.27.435072

**Authors:** Tomas Raul Wiche Salinas, Yuwei Zhang, Daniele Sarnello, Alexander Zhyvoloup, Laurence Raymond Marchand, Delphine Planas, Manivel Lodha, Debashree Chatterjee, Kasia Karwacz, Sally Oxenford, Jean-Pierre Routy, Heather Amrine-Madsen, Petronela Ancuta, Ariberto Fassati

**Affiliations:** Centre de recherche du Centre Hospitalier de l’Université de Montréal (CRCHUM), Université de Montréal, 900 Rue Saint-Denis, tour Viger, Montreal, Québec H2X 0A9, Canada; Institute of Immunity and Transplantation and Division of Infection & Immunity, University College London, the Rayne Building, 5 University Street, London WC1E 6JF, UK; Translational Research Office – Medicinal Chemistry, UCL School of Pharmacy, 29-39 Brunswick Square, London WC1N 1AX, UK; Division of Hematology and Chronic Viral Illness Service, McGill University Medical Centre (MUHC), 1001 Décarie Boulevard, Montreal, QC H4A 3J1, Canada; ViiV Healthcare, Five Moore Drive, Research Triangle Park, North Carolina, USA 27709-3398

## Abstract

Among CD4+ T-cells, T helper 17 (Th17) cells are particularly susceptible to HIV-1 infection and are depleted from mucosal sites, which causes damage to the gut barrier resulting in microbial translocation-induced systemic inflammation, a hallmark of disease progression. Furthermore, a proportion of latently infected Th17 cells persist long-term in the gastro-intestinal lymphatic tract, where low-level HIV-1 transcription is observed. This residual viremia contributes to chronic immune activation. Thus, Th17 cells are key players in HIV pathogenesis and viral persistence, however it is unclear why these cells are highly susceptible to HIV-1 infection. Th17 cell differentiation depends on expression of the master transcriptional regulator *RORC2, a* retinoic acid-related nuclear hormone receptor that regulates specific transcriptional programs by binding to promoter/enhancer DNA. Here, we report that RORC2 is a key host-cofactor for HIV replication in Th17 cells. We found that specific inhibitors that bind to the RORC2 ligand-binding domain reduced HIV replication in CD4+ T-cells. Depletion of RORC2 inhibited HIV-1 infection, whereas RORC2 overexpression enhanced it. RORC2 was found to promote HIV-1 gene expression. Chromatin immune precipitation revealed that RORC2 binds to the nuclear receptor responsive element (NRRE) in the HIV-1 LTR. In treated HIV-1 patients, RORC2+ CD4 T cells contained more proviral DNA than RORC2− cells. Pharmacological inhibition of RORC2 potently reduced HIV-1 outgrowth in CD4+ T-cells from antiretroviral-treated patients. Altogether, these results provide a new explanation as to why Th17 cells are highly susceptible to HIV-1 infection and point to RORC2 as a cell-specific target for HIV-1 therapy.

**Significance statement:** HIV-1 infects CD4 T cells and among these, Th17 cells are known to be particularly permissive for virus replication. Infection of Th17 cells is critical for AIDS pathogenesis and viral persistence, however it is not clear why these cells are highly permissive to HIV-1. We found that Th17 cell permissiveness depends on expression of the hormone receptor RORC2, which is the master transcriptional regulator of Th17 cell differentiation. We identify RORC2 as a new, cell-specific host-dependency factor that can be targeted by small molecules. Our results point to RORC2 as a cell-specific target for HIV-1 therapy, an entirely new concept in the field, and suggest HIV-1 might have evolved to exploit RORC2 to promote its own persistence.

## Introduction

A hallmark of HIV-1 infection is systemic inflammation, which can best predict disease progression (1, 2). A significant proportion of people living with HIV (PLWH) with undetectable plasma viral load during antiretroviral therapy (ART) have systemic inflammation, the severity of which correlates with overall mortality, morbidity and co-morbidity (1, 2) This inflammation also supports viral persistence because it promotes homeostatic proliferation and clonal expansion of memory CD4^+^ T cells carrying HIV-1 reservoirs (3, 4), enhances their migration into lymphatic organs and induces their activation, which stimulates local HIV-1 infection and reactivation from latency (5). The systemic inflammation observed in some virally suppressed individuals, indicates that widespread, ongoing virus replication is not the main cause; instead, substantial evidence suggests that inflammation has predominantly an indirect origin and is linked to alterations at mucosal sites (1, 6, 7).

T helper 17 (Th17) cells are a heterogeneous subset of CD4^+^ T-cells that expresses the chemokine receptor CCR6 and produce lineage-specific cytokines such as IL-17A, IL-21 and IL-22 (8–10). They are mainly found in the intestinal lamina propria and vaginal cervix mucosa, where they maintain the immunological barrier to microbiota, including bacteria and fungi (9–11). Remarkably, Th17 cells, which are preferentially targeted by HIV-1 *in vitro* and *in vivo* (10, 11) are also among the very first cells infected upon Simian immunodeficiency virus (SIV) vaginal exposure in macaques (12). As a consequence, Th17 cells are depleted from the gut and vaginal mucosa during acute infection in both PLWH (10, 11) and in SIV-infected monkeys (13, 14). The depletion of Th17 cells from the gastro intestinal lymphatic tract (GALT) of HIV-1 infected individuals has critical consequences for disease progression and viral persistence (15 Klatt, 2013 #255Dandekar, 2010 #529). Studies in PLWH and in pathogenic models of SIV infection showed that the loss of Th17 cells correlates with systemic inflammation, putatively via the disruption of the immunological homeostasis at the mucosal barriers and the translocation of bacterial products from the mucosa into the circulation (16, 17). Bacterial and fungal products then trigger the release of pro-inflammatory cytokines by various immune cells, establishing and maintaining systemic inflammation (7, 15, 17). These events are so critical that the loss or maintenance of Th17 cells can discriminate between pathogenic and non-pathogenic lentiviral infections (13, 15). Nevertheless, a proportion of infected Th17 cells are long-lived and constitute a component of the viral reservoir in the gut (8, 18–22), underscoring their importance in maintaining viral persistence during ART.

Despite these major insights, we do not fully understand why Th17 cells are preferentially targeted by HIV-1 for infection and how they are lost during the early acute phases of the disease (10, 11). Also, it is unclear which mechanisms govern HIV-1 reactivation from latency in long-lived Th17 cells carrying viral reservoirs in ART-treated PLWH.

The differentiation and effector functions of Th17 cells depend on expression of the master regulator retinoic acid receptor-related orphan receptor 2, RORC2 (RORγt in mice) (23–25). Here, we report that RORC2 is a bona fide host co-factor for HIV-1 gene expression in Th17 cells and identify small molecule RORC2 inhibitors that potently block HIV-1 replication/outgrowth without widespread cell toxicity. These findings establish a key molecular link between RORC2 expression and HIV-1 replication and point to RORC2 as a novel Th17 cell-specific target for HIV-1 therapy.

## Results

### RORC2 is a druggable host dependency factor for HIV-1

In a previous compound screening experiment to find small molecules with anti-HIV-1 activity, we identified two hits (digoxin and digitoxin) that are known antagonists of RORC2 (26, 27). This suggested that RORC2 may affect HIV-1 infection. To confirm and extend our initial observation, we tested several new, well-characterized RORC2 inhibitors developed by GlaxoSmithKline (GSK). Four of these compounds, GSK261805A (GSK805), GSK2837270A, GSK2793955A and GSK2837269A, were shown to bind to the RORC2 ligand-binding domain (LBD) with nanomolar affinity and displaced steroid receptor co-activator-1 (SRC1) (28) whereas GSK2833332A and GSK2805956A, had no measurable affinity up to 10 μM (**Table 1**). The four compounds with high affinity for RORC2 LBD also inhibited expression of a reporter luciferase gene under the control of the human IL-17A enhancer/promoter with sub-micromolar IC50s (**Table 1**), in agreement with the observation that RORC2 induces IL-17 expression by binding at promoter and enhancer regions in the IL-17 locus (29). Initially, we sought to test the compounds’ antiviral activity in Jurkat cells, whose susceptibility to HIV-1 infection is less variable than primary CD4^+^ T cells and does not require stimulation. To determine if Jurkat cells express RORC2 and hence may be sensitive to the compounds, we performed RT-qPCR using two sets of primers that specifically amplify either RORC2 or RORC1 (RORγ in mouse), an alternative spliced isoform that differs from RORC2 at its N-terminal region and is not normally expressed in CD4^+^ T cells (30). This analysis showed that Jurkat cells, similar to primary memory CD4^+^ T-cells, express RORC2 but not RORC1 transcripts (**Supporting information Figure 1a-b**); RORC2 protein expression was also confirmed in Jurkat cells by Western blot (**Supporting information Figure 1c**). A single cycle HIV-1 LAIΔenv virus expressing GFP and pseudotyped with VSV/G (hereafter called HIV-1 LAI_GFP_) (31) was used to infect Jurkat cells in the presence of increasing concentrations of RORC2 inhibitors or DMSO. Infected cells were analyzed by flow cytometry 48 hours post-infection and assessed for cell toxicity by alamarBlue™ or LIVE/DEAD staining. Compounds GSK2837269A and GSK2837270A inhibited HIV-1 LAI_GFP_ infection at a low micromolar range with no detectable cell toxicity, compound GSK2793955A showed intermediate potency whereas GSK2805956A and GSK2833332A showed lower potency **(Figure 1)**, in agreement with their low affinity for RORC2 (**Supporting information Table 1**).

**Table 1.** RORC2 compounds inhibit HTRF-ligand binding and Jurkat IL-17 luciferase reporter expression.

| **Compound** | **HTRF-Ligand Binding** |  | **Jurkat IL-17 Luciferase** |  |
| --- | --- | --- | --- | --- |
|  | **IC50 Mean μM (+/- SD), n** | **Vmax Mean (+/- SD), n** | **IC50 Mean μM (+/- SD), n** | **Vmax Mean (+/- SD), n** |
| GSK2691805A | 0.039 (0.001), 3 | 38 (10), 3 | 0.040 (0.002), 3 | 110 (0), 3 |
| GSK2837270A | 0.016 (<0.001), 2 | 94 (3), 2 | 0.501 (0.025), 2 | 110 (0), 2 |
| GSK2793955A | 0.025 (0), 1 | 86 (0), 1 | 0.794 (0.010), 2 | 110 (0), 2 |
| GSK2833332A | >10 (0), 1 | - | - | - |
| GSK2805956A | >10 (0), 1 | - | - | - |
| GSK2837269A | 0.016 (<0.001), 2 | 94 (4), 2 | 0.079 (0.001), 4 | 110 (1), 4 |

**Figure 1.**
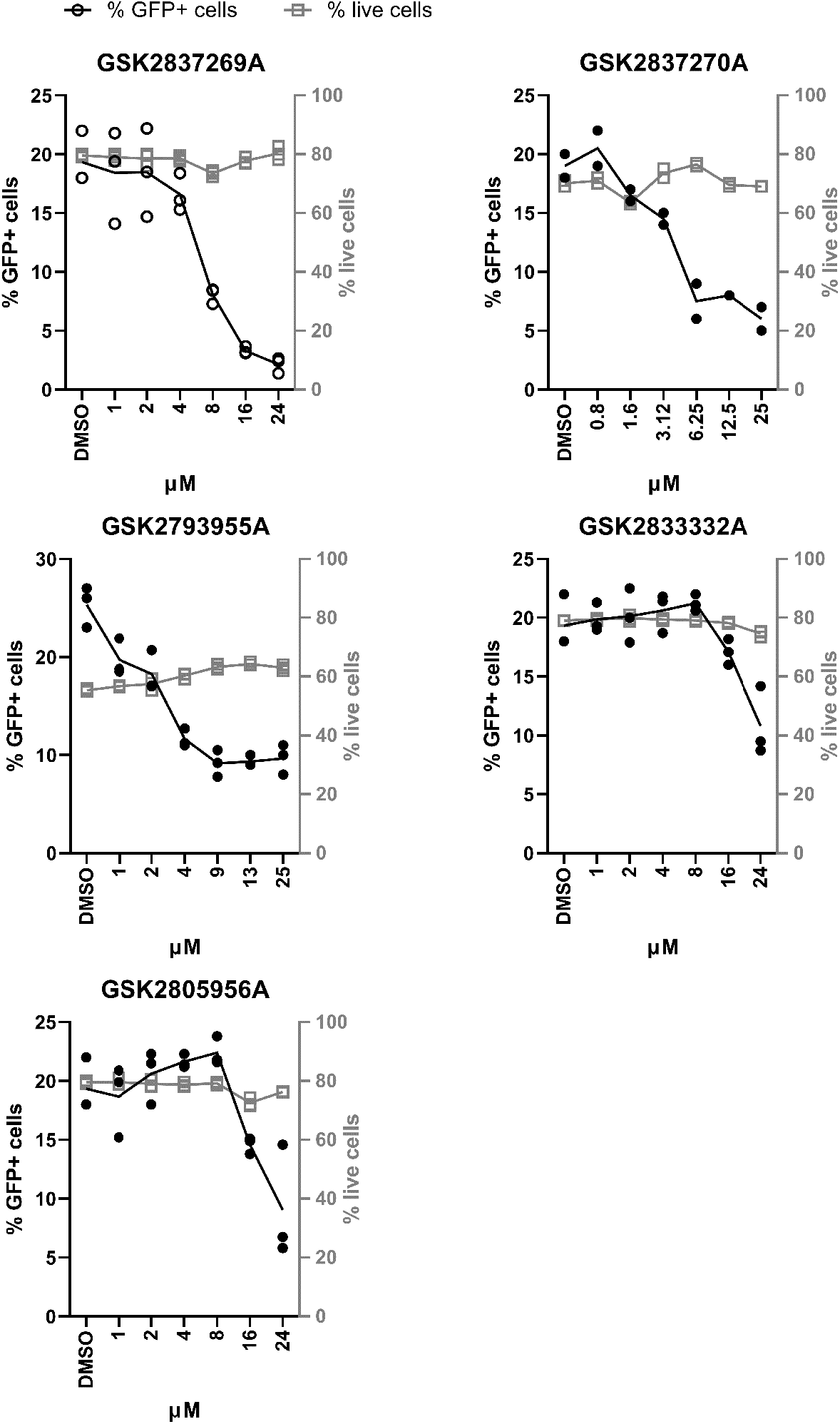
RORC inhibitors inhibit HIV-1 infection. Jurkat cells were infected with single cycle, VSVG-pseudotyped HIV-1 LAI_GFP_ at an MOI of 0.2 in the presence of the indicated concentrations of RORC2 inhibitors. Forty-eight hours post-infection, cells were analysed by flow cytometry to measure the percentage of GFP+ cells. The proportion of live cells was simultaneously assessed by AlamaR blue, n=3.

Next, we tested the activity of small molecule RORC2 inhibitors on HIV-1 replication in primary CD4^+^ T-cells. Memory CD4^+^ T cells were isolated from five HIV-uninfected healthy donors (HIV−) by magnetic sorting, stimulated for three days with CD3/CD28 antibodies (Abs) and infected with HIV-1_THRO_, a transmitted/founder virus (8), in the presence of the RORC2 inhibitors. Initially, we used GSK2691805A (also called GSK805), which is a prototypic RORC2 inhibitor structurally related to GSK2837269A (32, 33) (**Figure 2a)**. Infected cells were maintained in media containing IL-2 for 9 days after infection. IL-17A and interferon gamma (IFN-γ) were detected at day 3 post stimulation by ELISA. HIV-1 p24 released in cell culture supernatants was evaluated at day 3, 6 and 9 post- infection by ELISA; HIV-1 infected cells were detected by flow cytometry as CD4^low^HIV-p24^+^ at day 9 post-infection. We observed that GSK2691805A potently reduced HIV-1 replication, even at the lowest tested concentration of 0.3 μM (**Figure 2b-d**). RORC2 inhibitors GSK2837269A, GSK2793955A and GSK2833332A also reduced HIV-1 replication at a concentration of 5 μM, although differences did not reach statistical significance **(Figure 2e-f)**. We observed some differences in compounds’ potency between Jurkat and primary CD4^+^ T cells: GSK2837270A was active in Jurkat cells at 5 μM but not in primary CD4^+^ and conversely, GSK283332A was inactive in Jurkat cells but showed antiviral activity in primary CD4^+^ cells. These differences may be attributed to the different infection assays employed (single cycle in Jurkat cells vs replication over 9 days in primary cells), different intracellular concentrations of drugs due to distinct expression of drug transporters and efflux pumps (34), and possibly to different levels of RORC2 in these cells. All compounds, except for GSK2833332A, reduced IL-17A in a dose dependent manner but not IFN-γ production **(Supporting information Figure 2a-b**). This result suggested that GSK2833332A might inhibit HIV-1 infection in primary cells by a mechanism independent of RORC2. No cytotoxicity changes in cell proliferation for any of the compounds was observed at the tested concentrations (**Supporting information Figure 3a-b**). Using the same experimental design, GSK2691805A also reduced replication of HIV_NL4.3BaL_ (a molecular clone containing CCR5 tropic env obtained from the bronchoalveolar lavage (BaL) of an individual with chronic infection in a NL4.3 backbone) (data not shown). Therefore, inhibition of RORC2 impairs the replication of different HIV strains in Jurkat and primary CD4^+^ T- cell models.

**Figure 2.**
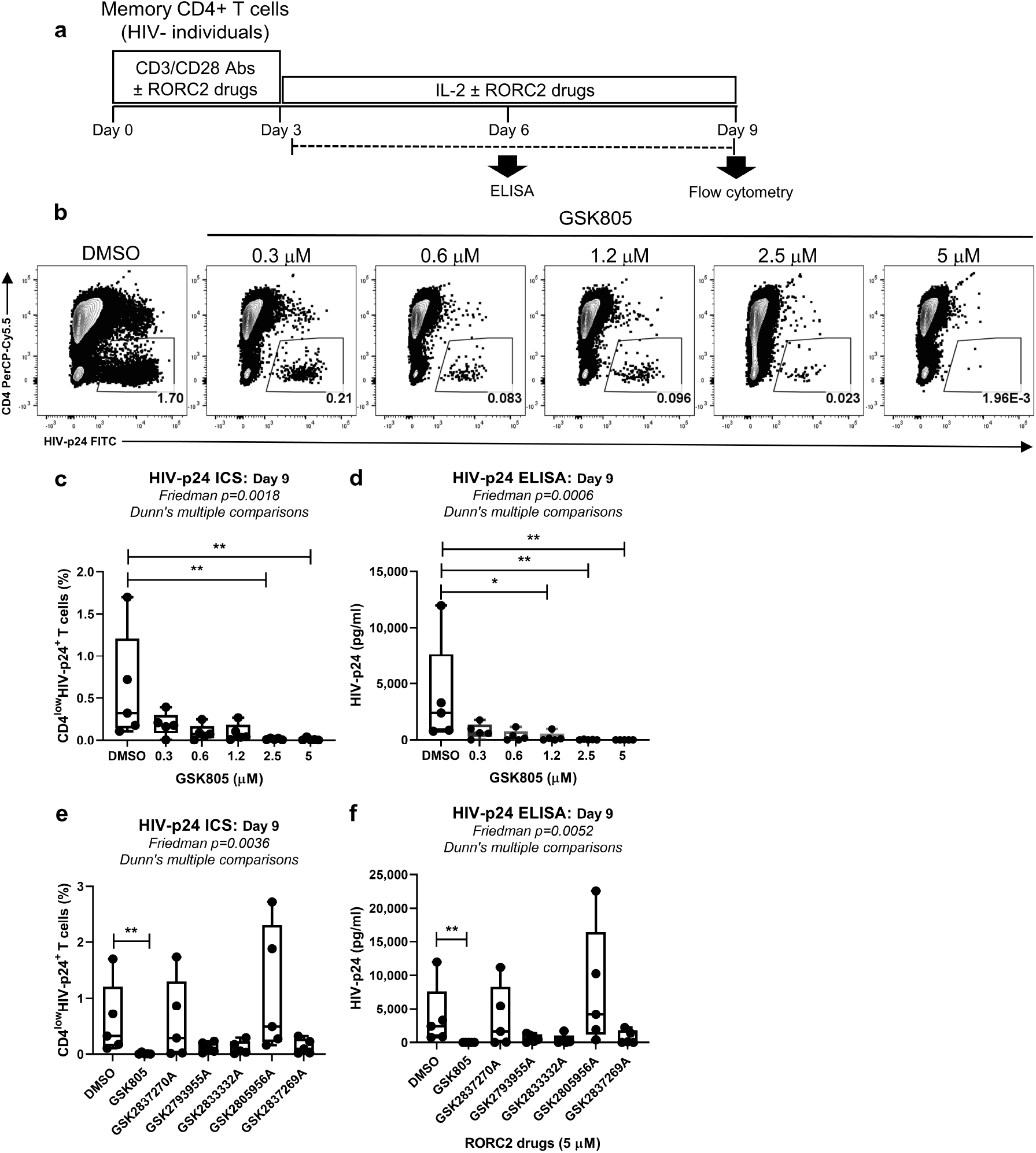
RORC inhibitors inhibit HIV-1 replication in primary CD4+ T cells. Shown is the experimental flow chart **(a)**. Memory CD4+ T cells isolated from n=5 healthy donors were stimulated with CD3/CD28 antibodies in the presence or absence of the indicated concentrations of RORC2 inhibitors for 3 days. Then, cells were exposed to HIVTHRO for 3 hours. HIV-infected CD4+ memory T cells were cultured with IL-2 in the presence of the indicated concentrations of RORC2 inhibitors for 9 days. Media was refreshed with IL-2 and RORC2 inhibitors every 3 days. **(b)** The intracellular expression of HIV-p24 in CD4+ memory T-cells was quantified by flow cytometry upon staining with fluorochrome-conjugated CD4 and HIV-p24 Abs at day 9 post-infection. Shown are contour plots of the frequency of CD4^low^HIV-p24^+^ T cells in one representative donor. **(c)** Statistical analysis of results obtained with cells from n=5 donors, when cells were cultured in the presence or the absence of GSK2691805A at the indicated concentration. **(d)** HIV-p24 levels measured in cell culture supernatants by ELISA. **(e)** Statistical analysis of the frequency of CD4^low^HIV-p24^+^ T cells and **(f)** HIV-p24 levels in cell-culture supernatants in the presence of the indicated RORC2 inhibitors (5 μM). Friedman p-values, with Dunn’s multiple comparison significance, are indicated on the graphs.

The pool of memory CD4+ T cells includes a Th17-polarized fraction, with the expression of Th17 effector functions in this fraction requiring T cell receptor (TCR) triggering (10, 35). To determine if RORC2 expression in this Th17-polarized population promoted HIV-1 infection *in vitro*, memory CD4^+^ T cells from healthy donors were stimulated for 3 days and infected with HIV-1_THRO_, (**Supporting information Figure 4a**). Six days post-infection, cells were analysed by flow cytometry to detect RORC2 and HIV-p24 expression (**Supporting information Figure 4b**). The percentage of HIV-p24+ cells was higher in RORC2+ than RORC2− cells in all five donors analysed, indicating better HIV-1 infection rates in cells expressing RORC2 (**Supporting information Figure 4c**). In parallel, CD4^+^ T cells from these same healthy donors were analysed by FACS to detect HIV-p24 expression in cells producing IL-17A and/or IFN-γ (IL-17A and IFN-γ are markers of Th17 cells and Th1 cells, respectively) upon simulation with PMA/Ionomycin (**Supporting information Figure 4d-f**). Notably, in these experimental settings, we confirmed that Th17 and Th17/Th1-polarized cells had a significantly higher proportion of HIV-p24+ cells than Th1 cells or unpolarised Th0 cells, consistent with their reported increased permissiveness to infection (10, 11).

To validate RORC2 as a host co-factor for HIV-1 infection, we generated three different hairpins targeting RORC2 mRNA and transduced them using lentiviral vectors into reporter Jurkat 1G5 cells, which express luciferase from the stably transfected HIV-1 LTR (36). After a short selection course to enrich for transduced cells expressing the shRNAs, we confirmed RORC2 knock-down (KD) by Western blot (**Figure 3a**) and infected the cells with HIV-1_NL4.3_. We found reduced HIV-1 replication in KD cells relative to control cells and this phenotype was proportional to the degree of RORC2 depletion, indicating specificity (**Figure 3b**). To test if RORC2 was sufficient to promote HIV-1 infection, we expressed RORC2 cDNA in 293T cells, which do not normally express endogenous RORC2 (**Figure 3c**), and infected them with HIV-1 LAI_GFP_ in the presence of DMSO or the RORC2 inhibitor GSK2837269A, which had the best activity/toxicity profile in Jurkat cells (**Figure 1**). Exogenous expression of RORC2 in 293T cells enhanced HIV-1 LAI_GFP_ infection above the already high basal levels and this phenotype was abrogated by GSK2837269A (**Figure 3d**). No inhibitory effect on infection was observed with GSK2837269A in control 293T cells at 12μM, consistent with the fact that they do not express the target. We also stably expressed a myc-tagged RORC2 cDNA in Jurkat cells using a retroviral vector construct and infected the cells with HIV-1 LAI_GFP_ virus. We found that Jurkat cells expressing exogenous RORC2 sustained better infection relative to control cells (**Figure 3e-f**), in agreement with the 293T cells data.

**Figure 3.**
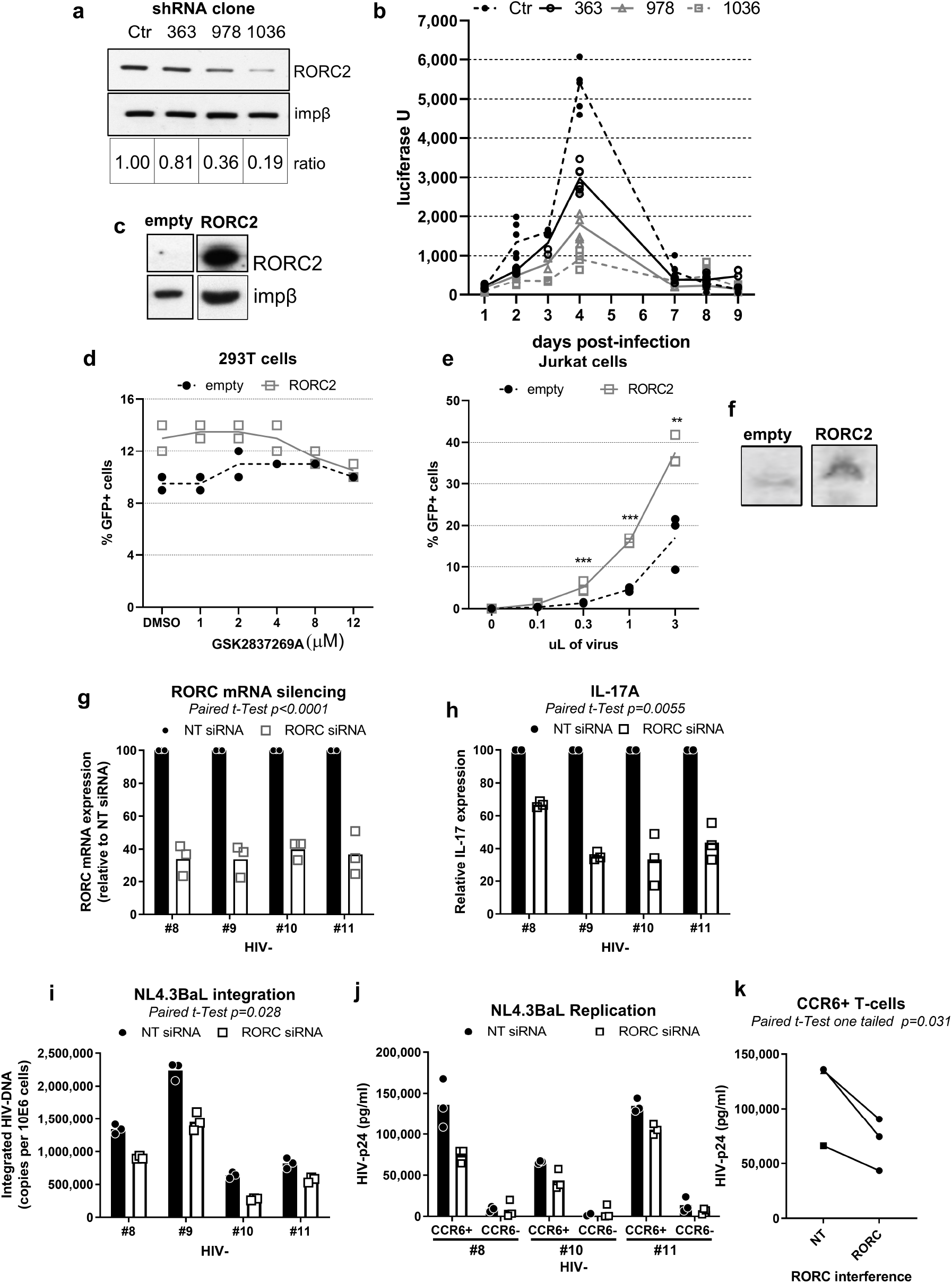
RORC2 is a host co-factor for HIV-1 infection of CD4+ T cells. **(a)** Luciferase Jurkat 1G5 indicator cells were transduced with three different shRNAs that target RORC2 or a non-targeting shRNA (Ctr) and selected in puromycin. RORC2 protein levels were analysed by Western blot and impβ was used as a loading control. The signal was quantified by ImageJ and the ratio of RORC2 versus impβ is indicated in the lower panel. **(b)** 1G5 cells transduced with the same shRNAs were infected with HIV-1_NL4.3_ and the luciferase signal was measured at the indicated time points. **(c)** An empty vector plasmid or the myc-RORC2 cDNA was transfected into 293T cells. After selection in puromycin, cells were analysed by Western blot as above. **(d)** The same 293T cells were infected with single cycle HIV-1 LAI_GFP_ at an MOI of 0.1 in the presence of the indicated concentrations of GSK2837269A and 40 hours later cells were analysed by flow cytometry to measure the percentage of GFP+ cells (n=2). **(e)** Jurkat cells were transduced with a retroviral vector expressing myc-RORC2 or an empty vector, selected in media containing blasticidin for 7 days and infected with the indicated volumes of single cycle HIV-1 LAI_GFP_ (10^7^ i.u./mL). Forty hours after infection cells were analysed by flow cytometry to measure the percentage of GFP+ cells. *** p<0.005, ** p = 0.008 two tailed Student’s T-test, n=3. **(f)** Expression of RORC2 in these cells was confirmed by Western blot. **(g-i)** Memory CD4+ T-cells from 4 healthy donors were stimulated *via* CD3/CD28 for 2 days and nucleofected with Dharmacon On target smart siRNA pools specific for RORC2 or a non-targeting (NT) siRNA using the Amaxa technology. **(g)** The efficiency of RORC2 silencing was assessed by real-time qPCR at day 1 post-nucleofection. **(h-i)** Cells were exposed to HIVNL4.3BaL (50 ng per 10^6^ cells) then IL-17A and HIV-DNA integration were quantified by ELISA in cell-culture supernatants and nested real-time PCR in cell pellets, respectively, at day 3 post-infection. **(j-k)** RORC2 RNA interference was performed on sorted memory CCR6+ and CCR6− CD4+ T-cells from n=3 individuals, infected as described above and analysed 3 days post-infection to determine the levels of HIV-p24 in cell-culture supernatants **(j).** Statistical analysis of HIV-p24 levels in CCR6+ T-cells sorted from n=3 individuals **(k)**.

We next sought to validate RORC2 as a co-factor for HIV-1 infection by mRNA depletion in primary CD4 T cells. Memory CD4 T cells were CD3/CD28-stimulated, nucleofected the next day with a siRNA targeting RORC2 mRNA or a non-targeting control siRNA and infected with HIV-1_BaL_ one day later. We observed partial inhibition of the RORC2 mRNA expression in all the tested donors (**Figure 3g**), which resulted in lower IL-17A production from the treated cells relative to controls (**Figure 3h),** and a small but significant reduction of HIV-1 infection as measured by the quantity of integrated viral DNA accumulated over the 3 days period (**Figure 3i**). Finally, because RORC2+ Th17 cells express CCR6 (37), RORC2 siRNA experiments were similarly performed on flow cytometry-sorted CCR6+ and CCR6-CD4+ T cells. The highest levels of HIV replication were detected in CCR6+ T cells, in agreement with previous reports by our group and others (8, 18, 19, 21, 22). Noteworthy, depletion of RORC2 mRNA resulted in a significant reduction in integrated viral DNA and HIV-p24 levels in CCR6+ subset, with no effects observed in CCR6-T cells, indicative of decreased HIV-1 infection/replication upon RORC2 depletion (**Figure 3j-k).** Taken together, these pharmacological and genetic approaches demonstrated that RORC2 promote HIV-1 replication.

### RORC2 regulates HIV-1 gene expression

To determine the step of the viral replication cycle that was impaired by the anti-RORC2 compounds, we infected Jurkat cells with HIV-1 LAI_GFP_ at an MOI of 0.1 in the presence of GSK2837269A, GSK2837270A, or nevirapine (a non-nucleoside inhibitor of reverse transcriptase), raltegravir (a strand transfer inhibitor of integration) or DMSO. Total DNA was extracted from the cells at 24 hours post-infection and Taqman qPCR was used to measure the amount of GFP DNA (as a surrogate marker of negative strand viral DNA) and 2LTRs circular DNA, a hallmark of nuclear entry (38). To measure the amount of integrated viral DNA, we performed Alu-LTR quantitative PCR from DNA extracted 8 days post-infection (27). RORC2 inhibition did not significantly impair reverse transcription, nuclear entry or integration, in contrast to nevirapine, which reduced both viral DNA and 2LTRs, or raltegravir, which suppressed integration (**Figure 4a**). These results indicated that RORC2 acts at a step post integration, such as gene expression.

**Figure 4.**
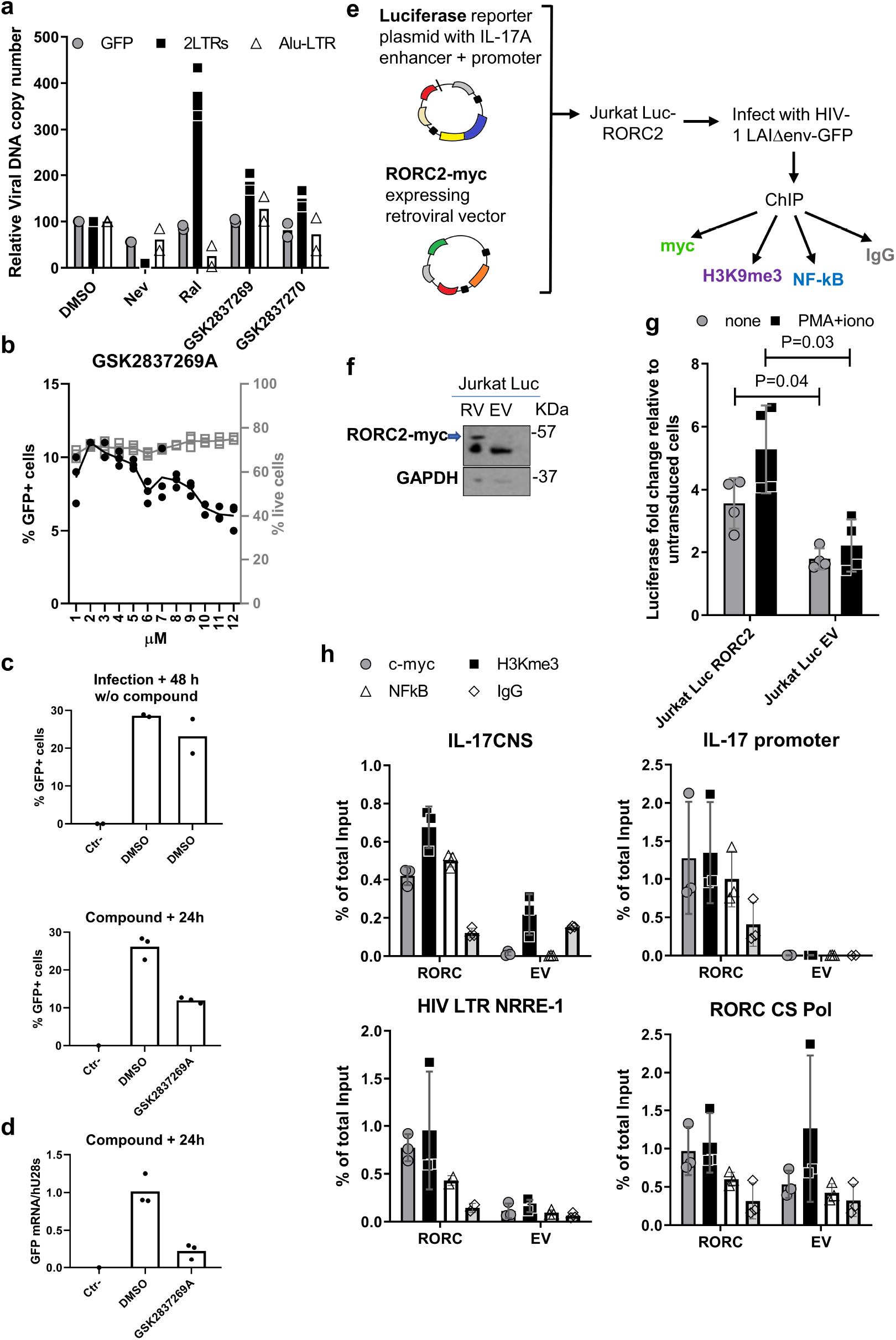
RORC2 promotes HIV-1 gene expression and binds to the HIV-1 LTR. **(a)** Jurkat cells were infected with VSV-G pseudotyped single cycle HIV-1 LAI_GFP_ in the presence of the reverse transcriptase inhibitor nevirapine (500 nM), or the integrase inhibitor raltergravir (100 nM) or the indicated RORC inhibitors (5 μM). The amount of negative strand viral DNA (GFP) and 2-LTR circular DNA was quantified 24h post-infection by Taqman qPCR. Integrated viral DNA was quantified by Alu-LTR qPCR on DNA extracted 8 days after infection, n=2. **(b)** Jurkat cells were infected with VSV-G pseudotyped single cycle HIV-1 LAI_GFP_ and 24h post-infection they were treated with the indicated concentrations of GSK2837269A. Cells were analysed 24h later by flow cytometry to determine the percentage of infected (GFP+) cells, n=3. **(c)** Jurkat cells were infected with single cycle HIV-1 LAI_GFP_. Forty-hours after infection, cells were analysed by flow cytometry to determine the percentage of infected (GFP+) cells. These cells were then treated with DMSO or 5 μM GSK2837269A for 24h before re-analysis by flow cytometry. **(d)** RNA was extracted from the cells and the amount of GFP mRNA relative to 28S ribosomal RNA was quantified by RT-qPCR. Average values are shown (n=3). **(e)** Schematic diagram explaining the experimental steps for ChIP. Jurkat cells expressing luciferase driven by the IL-17A enhancer/promoter region were transduced with a retroviral vector expressing RORC2- myc (Luc -RORC2) or an empty vector control (Luc-EV). Luc-RORC2 and Luc-EV cells were infected in parallel with VSV-G pseudotyped HIV-1 LAI ΔenvGFP (HIV-1GFP) and 24 hours later processed for ChIP with the four indicated antibodies. **(f)** Western blot with anti-myc antibody to detect RORC-myc following ChIP (the lower band is IgG heavy chains); GAPDH in the total lysate samples was used as a loading control. **(g)** Fold change in Luciferase expression relative to untransduced Jurkat Luc cells with or without stimulation with PMA and ionomycin. Shown is average ± SD, n=3, P value based on two tailed t-test. **(h)** Real time PCR signal following ChIP for the indicated DNA regions and antibody. Top left, IL-17A enhancer region; top right, IL-17A promoter region; bottom left, NRRE region in the HIV-1 LTR; bottom right, HIV-1 pol region containing the putative RORC2 consensus element. Shown are replicate values, as well as average ± SD, n=3.

To examine this aspect further, GSK2837269A was added on to Jurkat cells chronically infected with HIV-1 LAI_GFP_ and the percentage of GFP+ cells was analysed by flow cytometry 48 hours later. A dose-dependent reduction of the percentage of GFP+ cells was observed (**Figure 4b**). To examine viral gene expression directly, Jurkat cells were infected with HIV-1 LAI_GFP_ in the absence of compounds and analysed by flow cytometry 48 hours later (**Figure 4c top panel**). Next, GSK2837269A was added onto these infected cells, which were reanalysed by flow cytometry 24 hours later (**Figure 4c middle panel**). In parallel, RNA was extracted from the GSK2837269A-treated cells and used for RT-qPCR to detect GFP mRNA, which is transcribed from the viral LTR. At 5 μM, GSK2837269A reduced GFP mRNA 5-fold (**Figure 4d)**. The modest discrepancy between GFP mRNA and percentage of GFP+ cells is most likely explained by the longer half-life of the GFP protein relative to its mRNA.

RORC2 binds to specific DNA consensus elements, recruiting chromatin-activating co-factors via its ligand-binding domain to regulate transcription (23, 39). We therefore hypothesized that RORC2 might bind to such DNA elements on the HIV-1 provirus. The HIV-1 LTR contains a well characterized nuclear receptor responsive element (NRRE) that binds several retinoic acid receptors (40, 41). In addition, we have detected *in silico* a consensus RORC DNA binding motif (23, 32) in the pol region of the HIV-1 genome (**Supporting information Figure 5).** It has previously been reported that this region in Pol may have enhancer activity (42). To test the hypothesis, we performed chromatin immunoprecipitation followed by real-time PCR (ChIP-qPCR) for both the NRRE in the LTR and the RORC2 consensus sequence in Pol (CS Pol). There is no available ChIP-grade antibody against human RORC2 hence we performed the experiments in Jurkat cells stably transduced with a retroviral vector expressing C-terminally myc-tagged RORC2. However, Jurkat cells do not express IL-17A, raising the possibility that the IL-17A locus might be defective thus making these cells unsuitable to detect RORC2 binding to the IL-17A regulatory elements by ChIP. To circumvent this problem, we transduced with the RORC2-myc expressing retroviral vector Jurkat cells stably transfected with a plasmid expressing luciferase driven by the IL-17A promoter and CNS-5 enhancer, which contain one RORC2 DNA consensus element each (Jurkat-Luc cells) (29, 43) (**Figure 4e-f**). Jurkat-Luc cells transduced with an empty retroviral vector (Jurkat Luc-EV) were generated to control for specificity. The resulting Jurkat Luc-RORC2 cells expressed luciferase at higher levels than Jurkat Luc-EV both at baseline and after stimulation, confirming their functionality (**Figure 4g**). Jurkat Luc-RORC2 and Luc-EV cells were infected in parallel experiments with HIV-1 LAI_GFP_ at low MOI. Twenty-four hours after infection, cells were processed for ChIP using an anti-myc antibody. We also employed an antibody against H3K9me3, which is a histone marker of inactive but poised enhancer/promoters (44) and an antibody against NF-κB, a transcription factor known to bind both the HIV-1 LTR (45) and, in certain conditions, the IL-17 enhancer/promoter regions (46, 47). A specific ChIP signal for RORC2-myc was detected on the IL-17 enhancer and promoter regions of Jurkat Luc-RORC2 cells but not Jurkat Luc-EV cells (**Figure 4h**). Notably, a specific ChIP-qPCR signal was observed on the HIV LTR NRRE element whereas the signal for the CS Pol element was less convincing due to high background, which may be related to greater “stickiness” of the specific DNA sequence under study due to charge and/or secondary structure (**Figure 4h**).

These results showed that RORC2 binds to the HIV-1 LTR with similar or even greater strength than the IL-17 promoter/enhancer regions. Notably, a specific ChIP-qPCR signal was also detected for H3K9me3 in Jurkat-Luc cells, particularly on the IL-17A enhancer/promoter region. This H3K9me3 signal was stronger in RORC2-expressing cells, suggesting that RORC2 might either associate with, or increase the proportion of poised enhancers. It should be noted that Jurkat cells were not stimulated before ChIP, hence some critical RORC2 co-factors important for chromatin remodelling might be present at low levels (23, 29, 47), reducing the rate of enhancer conversion from poised into active. Furthermore, the RORC2− specific ChIP-qPCR signal is likely to be an underestimation due to competition with the endogenous RORC2, which is expressed in Jurkat cells but cannot be precipitated by the anti-myc Ab (**Supporting information Figure 1**). Consistent with previous reports, we did not detect NF-κB binding to the IL-17 enhance/promoter in the absence of stimulation (46, 47), however expression of RORC2 appeared to stimulate recruitment of NF-kB on both the IL-17 enhancer/promoter region and the HIV-1 LTR, in agreement with the observed co-operativity of RORC2 and some transcription factors (23, 29, 46). Taken together, these results support the possibility that RORC2 regulates viral gene expression by binding to the HIV-1 LTR.

### RORC2 is critical for HIV-1 outgrowth from patients’ cells

Since RORC2 promotes HIV-1 gene expression in acutely infected cells, we tested if its inhibition also prevented HIV-1 outgrowth from cells of PLWH. Since RORC2-mediated effector functions are not constitutive in Th17 cells but dependent on TCR triggering (10, 35), in preliminary experiments we evaluated if the activation of memory CD4^+^ T-cells with CD3/CD28 antibodies was capable of inducing the expression of RORC2 at the transcriptional and protein level. Memory CD4^+^ T-cells were isolated from HIV-uninfected individuals, stimulated for 5 or 24h hours. The relative RORC2 mRNA and protein levels were assessed 5 or 24 hours later, respectively. We observed that cell activation induced the expression of RORC2 but not RORC1 mRNA **(Supporting information Figure 6a)**. At the mRNA and protein level, RORC2 expression was up-regulated by TCR activation and cells expressing RORC2 also expressed CCR6 **(Supporting information Figure 6b-d**), which is a well-established marker of human Th17 cells (9) and HIV-reservoir enrichment (8, 10, 18, 22, 48). Next, to determine if cells expressing RORC2 harbour HIV-1, memory CD4 T cells were isolated from ART+PLWH and stimulated *in vitro* with anti-CD3/CD28 Abs for 24 hours in the presence of ARVs to prevent HIV cell-to-cell transmission *in vitro* **(Figure 5a)**. This stimulation is required for optimal expression of RORC2, which, similar to all lineage-specific cytokines, is not constitutively expressed in Th17-committed CCR6+ T cells (35). Cells were then sorted based on CCR6 and/or RORC2 expression **(Figure 5b)** and analysed by PCR to detect integrated HIV DNA **(Figure 5c**). In all five ART+PLWH, proviral DNA was significantly more abundant in CCR6+RORC2+ cells compared to CCR6−RORC2− cells or CCR6+RORC2− cells (**Figure 5c),** indicative of preferential infection in CCR6+RORC2+ cells. To explore whether CCR6+RORC2+ cells carry replication-competent HIV-DNA, we performed a modified HIV Flow Assay to quantify HIV-p24 expression in RORC2+ cells (49). To ensure detection of infected cells, studies were performed with cells from viremic untreated PLWH (ART− PLWH). To this end, memory CD4^+^ T-cells isolated from ART− PLWH were stimulated for 72 hours in the presence of ARVs and RORC2 expression measured in total T cells and productively infected CD4^low^HIV-p24+ T cells. As expected, we found that cell stimulation induced the expression of both RORC2 and HIV-p24 (data not shown). Notably, CD4^low^HIV-p24+ T cells were enriched in RORC2 expression compared to total memory T cells **(Supporting information Figure 7a-c)**. Additionally, CD4^low^HIV-p24+ T cells expressing RORC2 showed a higher HIV-p24 GMFI (geometric mean fluorescence intensity) compared to their RORC2− counterparts **(Supporting information Figure 7d)**, indicating that a fraction of RORC2+ T cells harbour translation-competent proviruses.

**Figure 5.**
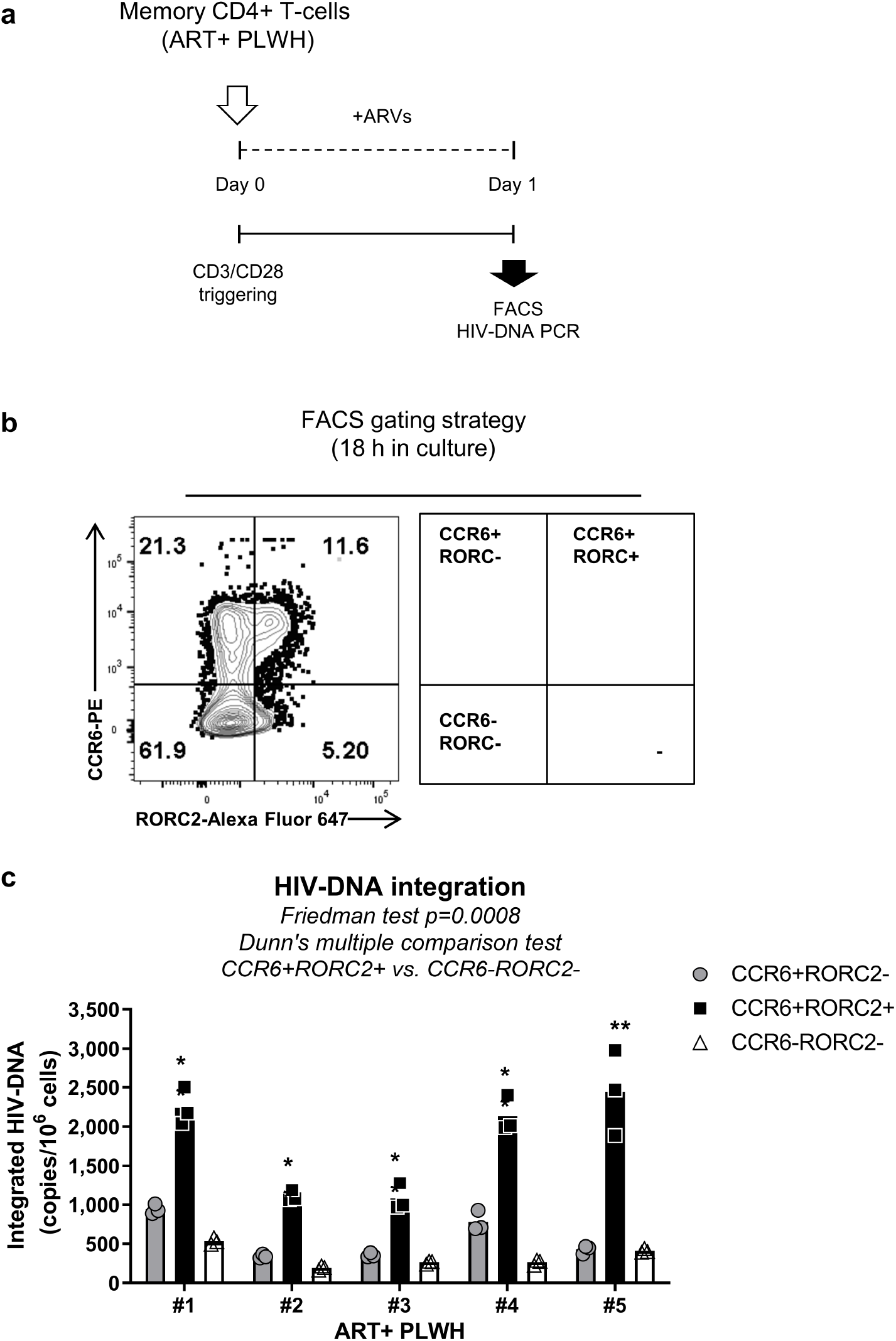
Memory CD4+ T-cells expressing the Th17 markers CCR6 and RORC2 are enriched in integrated HIV-DNA in ART-treated PLWH. **(a)** Flow chart of experimental approach; memory CD4^+^ T-cells isolated from ART+ PLWH were stimulated with CD3/CD28 Abs for 18 hrs in the presence of antiretroviral drugs (AZT 180 nM, Efavirens 100nM, Raltegravir 200nM) to prevent cell-to-cell HIV transmission *in vitro*. Highly pure CCR6+RORC2−, CCR6+RORC2+ and CCR6−RORC2− cell subsets were sorted by FACS and Integrated HIV-DNA levels were quantified by nested real-time PCR. **(b)** Gating strategy used to sort the three cell populations mentioned above and **(c)** the statistical analysis of Integrated HIV-DNA in sorted CD4+ T-cell subsets from n=5 ART-treated PLWH. Shown are individual replicates, with the bars representing median values.

Based on these observations, we sought to test if RORC2 inhibition prevented HIV-1 reactivation from latency and/or viral outgrowth *ex vivo*. To this end, we performed a simplified viral outgrowth assay (VOA) we have previously described (50) using memory CD4^+^ T cells from both ART+ and ART− PLWH. Cells were stimulated with anti-CD3/CD28 Abs for 3 days and maintained in culture for another 9 days by splitting each well into two new wells every 3 days, in the presence of DMSO or 5 μM GSK2691805A (**Figure 6a**). Cells were analysed to detect CD4^low^HIV-p24+ cells by flow cytometry at day 12 post CD3/CD28 antibody stimulation and the levels of HIV-p24 in cell-culture supernatants were evaluated at day 9 and 12 post-activation by ELISA. In the presence of GSK2691805A, there was a consistent reduction in the frequency of CD4^low^HIV-p24+ cells compared to DMSO on T cells from both ART+ and ART− PLWH (**Figure 6b-d**). Similarly, HIV-p24 levels in the culture supernatants were significantly reduced by GSK2691805A treatment (**Figure 6e-f**). These results are consistent with the notion that RORC2 is critical for HIV-1 reactivation/outgrowth in infected Th17 cells from PLWH.

**Figure 6.**
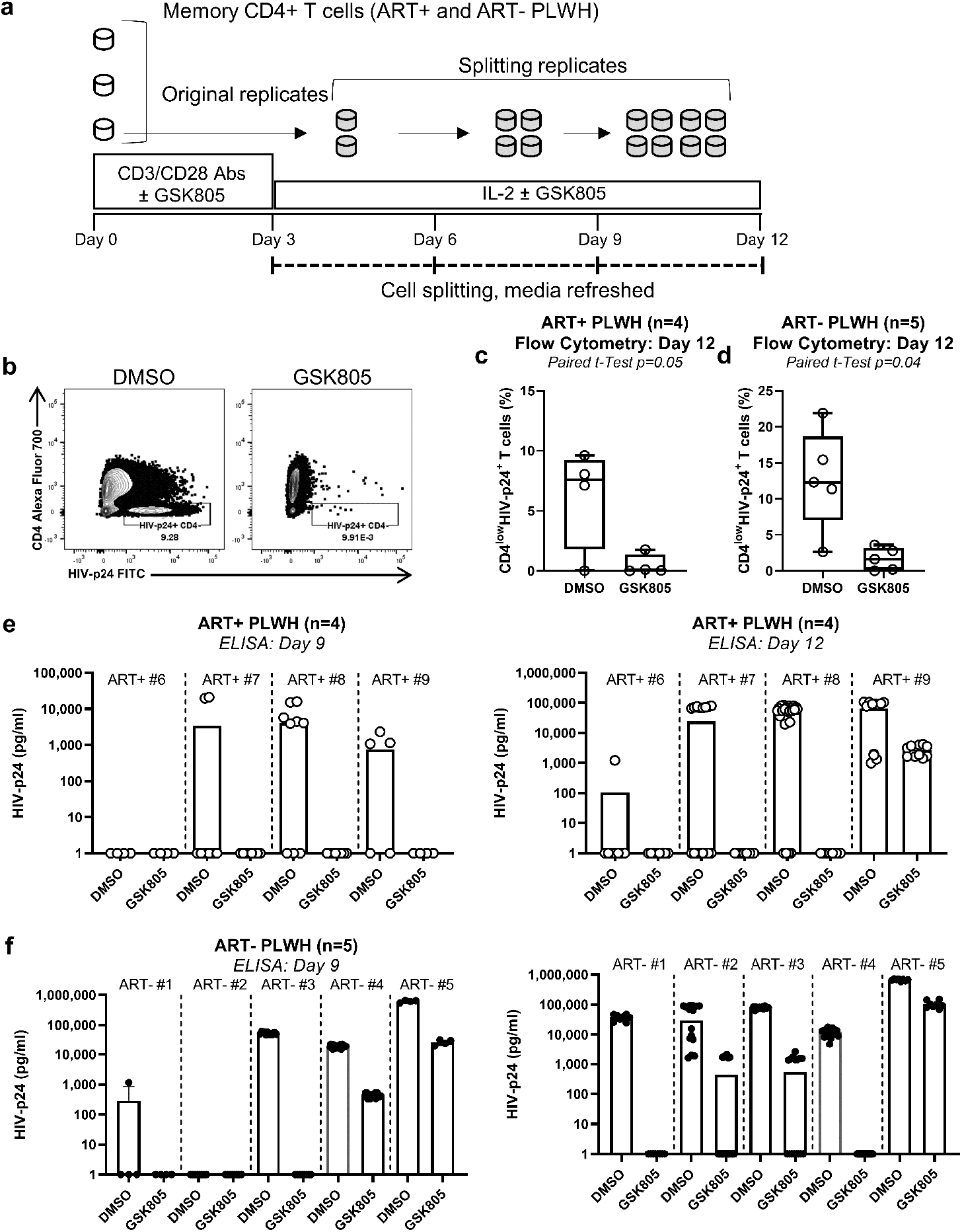
RORC antagonism inhibits HIV-1 outgrowth in memory CD4+ T cells of ART-treated and untreated PLWH. **(a)** Experimental flow chart of the viral outgrowth assay; memory CD4+ T-cells isolated from PLWH receiving ART (ART+PLWH) or not (ART− PLWH) were activated with CD3/CD28 Abs in the presence of DMSO or GSK2691805A (5 μM) at a cell concentration of 1 million cells/ml in triplicates for 3 days. Then, cells were washed and split at day 3 post-stimulation and cultured in medium containing IL-2 (5 ng/ml) up to 12 days (split every three days) in the presence of DMSO or GSK2691805A (5 μM). At day 12, HIV-infected cells were identified as CD4^low^HIV-p24+ by flow cytometry. HIV-p24 levels in cell-culture supernatants were measured by ELISA. **(b)** Frequency of CD4^low^HIV-p24+ cells in one representative individual. **(c-d)** Statistical analysis of results obtained with cells of n=4 ART+PLWH **(c)** and n=5 ART-PLWH **(d)**. **(e-f)** Effect of GSK2691805A on HIV-p24 levels in cell-culture supernatants of n=4 ART+PLWH **(e)** and n=5 ART-PLWH **(f)** at day 9 and 12 post CD3/CD28 activation. Paired t-Test p-values are indicated on the graphs. Shown are box and whiskers plots with individual values and maximum and minimum **(c-d),** and individual replicate values, with bars indicating median values **(e-f)**.

## Discussion

In this study, we have identified the Th17 cell master transcription regulator RORC2 as a host co-factor for HIV-1 infection and showed that it regulates viral gene expression upon infection *in vitro* and possibly reactivation from latency, as reflected by viral outgrowth *ex vivo*. These findings provide a new explanation as to why HIV-1 replicates more efficiently in Th17 cells, which express higher levels of RORC2 relative to other CD4^+^ T cell types. Current antiretroviral drugs inhibit various steps of the HIV-1 replication cycle but not transcription. Thus, RORC2 inhibitors represent a new class of anti-HIV molecules that may limit the deleterious consequences of HIV-mediated Th17 cell depletion from mucosal sites before and after ART initiation.

It is now well-established that a proportion of latently infected Th17 cells do persist long-term (8, 20, 22) and constitute a sizeable fraction of the viral reservoir in the GALT (18, 21, 48), a tissue where persistent low-level HIV-1 transcription is observed (51, 52). This residual transcription contributes to chronic immune activation/exhaustion leading to serious non-AIDS co-morbidities (51). Noteworthy, such transcriptionally active viral reservoir appears to be the source of viral rebound upon ART interruption (53). Because RORC2 inhibitors repress HIV-1 gene expression, these molecules may also be used in “block and lock” strategies to counteract residual transcription in long-lived Th17 cells carrying viral reservoirs during ART and to prevent viral rebound upon treatment interruption.

The ChIP-qPCR results indicate that RORC2 binds to the NRRE in the HIV-1 LTR. This DNA element is well conserved among different HIV-1 subtypes and was previously shown to be recognized by several nuclear hormone receptors, including retinoic acid receptors that positively regulate viral gene expression in CCR6+ Th17 cells (8, 11, 18). It is reasonable to propose that RORC2 has a similar mechanism of action. The NRRE in the HIV-1 LTR might be a critical element providing transcriptional plasticity to the virus resulting in better adaptation to different cell types (54). It is noteworthy that oestrogen inhibits HIV-1 reactivation from latency in Th17 cells (55). This effect is mediated by binding of the nuclear hormone receptor oestrogen receptor-1 (ESR-1) to the LTR, which may explain why women appear to have a lower inducible reservoir than men (55). Oestrogen also inhibits Th17 cell differentiation and IL-17 secretion (56, 57), suggesting a broader interconnection between oestrogen and RORC2. It will be interesting to test if RORC2 and ESR-1 compete for binding to the LTR and exert antagonistic effects on viral transcription.

We observed that RORC2 expression increased the fraction of the NRRE that was precipitated by an antibody against histone H3K9me3, a marker of inactive but poised enhancers (44). The same was found with the IL-17 enhancer/promoter, suggesting that it was not LTR-specific. The reasons for this are unclear but we speculate that RORC2, when additional co-factors are rate-limiting, may induce poising of specific enhancers but alone may be insufficient to fully activate them. This is consistent with the notion that co-operation is necessary between RORC2 and transcription factors such as BATF, IRF4, STAT3 and Runx1 to activate the transcriptional program leading to Th17 cell differentiation (23, 29, 43, 46).

We therefore propose that RORC2, rather than being essential for HIV-1 gene expression, acts more like a positive modulator of HIV-1 transcription, perhaps by helping establish a suitable chromatin environment. This may explain why Th17 cells, which express high levels of RORC2, sustain better HIV-1 replication. Higher HIV-1 gene and protein expression in Th17 cells may also make these cells better targets for killing by CD8^+^ T cells and in part explain long-term depletion of such cells from the GALT. In addition to a direct role in HIV-1 gene expression, RORC2 may have other indirect effects that promote viral replication via the transcriptional regulation of other HIV-1 permissiveness factors (58) and these effects are worthy of further research. It will be interesting to investigate if HIV has evolved to recruit RORC2 to boost viral gene expression and increase its replication efficiency in Th17 cells, a subset enriched in frequency at portal sites of viral entry (10, 11). This may provide an evolutionary advantage for the initial steps of viral transmission, with the loss of Th17 cells leading to systemic inflammation and CD4^+^ T cells activation (10, 11, 59), creating a suitable environment for virus persistence.

Our results demonstrate that the pharmacological inhibition of RORC2 potently suppressed HIV-1 outgrowth *ex vivo* in cells from ART-treated or untreated PLWH. This is consistent with the idea that RORC2 stimulates HIV-1 gene expression and may be important in regulating the dynamics of the viral reservoir (60). Fluctuations in RORC2 expression levels correlate with the activation status of Th17 cells, which is dependent, among other things, on priming via the TCR by specific antigens from pathogens such as *Candida albicans* and *Staphylococcus aureus* (9). Thus, specific stimuli from the microbiota may upregulate RORC2 expression selectively in latently infected Th17 cells, promoting reactivation from latency and viral rebound.

Pharmacological inhibition of RORC2 *in vivo* may help elucidate the contribution of Th17 cells on the latent viral reservoir. The GSK RORC2 inhibitors have been employed in animal studies for non-HIV indications and GSK2691805A has been used to study the effect of RORC2 inhibition on Th17 and ILC-3 cells in mice (61). It will therefore be possible to conduct experiments in pre-clinical animal models of HIV-1 infection to determine if RORC2 inhibition delays or prevents viral rebound upon ART interruption and/or whether RORC2 inhibition reduces inflammation and immune activation. A limitation of this approach is that Th17 cell differentiation and their effector functions might also be affected by administration of the RORC2 inhibitors. Nevertheless, the effects of RORC2 drugs may be reversible, as indicated by the capacity of cells to produce IL-17 again upon drug withdrawal *in vitro* (data not shown). Optimal dosing regimens and regular monitoring of Th17 cell populations in various tissues may allow the safe testing of RORC2 inhibitors in pre-clinical and clinical interventions. Other potential limitations are linked to the fact that other cells also express RORC2, including ILC-3 and thymocytes (62, 63), although ILC-3 seem tolerant to prolonged inhibition of RORC2 (61). These RORC2+ cells will need to be carefully monitored too. It might also be possible to use RORC2 inhibitors for short-term interventions in the early stages of infection or longer-term at low doses to inhibit residual HIV-1 transcription/translation in Th17 cells, in an effort to minimise inflammation and immune activation documented in ART-treated PLWH. Despite these potential limitations, which may be mitigated, targeting RORC2 *in vivo* is a conceptually new approach that affords a unique opportunity to reduce HIV-1 replication and reactivation from latency in a specific cell population highly relevant for disease progression.

## Materials and Methods

### Ethics statement

The collection of leukapheresis from HIV-uninfected individuals, ART− PLWH and ART+ PLWH was conducted in compliance with the principles included in the Declaration of Helsinki. This study received approval from the Institutional Review Board of the McGill University Health Centre and the CHUM-Research Centre, Montreal, Quebec, Canada. Written informed consents were obtained from all study participants.

### Human subjects

The human biological samples were sourced ethically and their research use was in accord with the terms of the informed consents under an IRB/EC approved protocol. HIV-uninfected individuals [HIV−; n= 18; 17 males and 1 female; median age of 57.5 years (range: 25-70), median CD4 counts 752 cells/μl (range: 511-1,115)], as well as virally suppressed ART-treated PLWH [ART+ PLWH; n=9; 9 males and 0 female; median age of 44 years (range: 30-57), median CD4 counts 514 cells/μl (range: 318-598)] and untreated PLWH (ART− PLWH; n=6; 6 males and 0 female; median age of 41 years (range: 24-50), median CD4 counts 459 cells/μl (range: 221-1,068)]) (**Supporting information Table 2**), were recruited at the Montreal Chest Institute, McGill University Health Centre and Centre Hospitalier de l’Université de Montréal (CHUM, Montreal, QC, Canada). Peripheral blood mononuclear cells (PBMC) (10^9^−10^10^ cells) were collected by leukapheresis and frozen until use, as previously described (8, 18, 20). Plasma viral load in ART-treated PLWH was measured using the Amplicor HIV-1 monitor ultrasensitive method (Roche).

### HIV infection *in vitro* of Primary memory CD4+ T-cell

The molecular clones of CCR5-tropic HIV-1 strain used in this study were: transmitted Founder (T/F) THRO and NL4.3BAL HIV-1. The T/F THRO HIV plasmid was obtained through the NIH AIDS Reagent Program, Division of AIDS, NIAID, NIH: pTHRO.c/2626 (cat# 11745) from Dr. John Kappes and Dr. Christina Ochsenbauer. The NL4.3BaL HIV plasmid was provided by Dr. Michel Tremblay (Univerité Laval, Québec, Canada), originating from Dr Roger J Pomerantz (Thomas Jefferson University, Philadelphia, Pennsylvania, USA). HIV-1 plasmid was amplified by MiniPrep and MaxiPrep and viral stocks were produced and titrated, as we described it previously (8, 20). Memory CD4^+^ T-cells were isolated from PBMCs of HIV-uninfected individuals by negative selection using magnetic beads (Miltenyi Biotec, Auburn, CA, USA), as we previously described (8, 20). Then, memory CD4^+^ T-cells (1×10^6^ cells/ml per 48 well-plate well) were stimulated with immobilized CD3 and soluble CD28 Abs (1 μg/mL) for 3 days prior to infection. T-cells were cultured with RPMI1640 media (Gibco, Grand Island, NY, USA) containing 10% FBS and 1% P/S. Memory CD4^+^ T-cells were infected with HIV-1 (50 ng HIV-p24/10^6^ cells) and then cultured in the presence of IL-2 (5 ng/ml; R&D Systems, Minneapolis, MN, USA) for up to 9 days, with 50% of the media being refreshed every 3 days. Viral replication was measured by flow cytometry analysis upon intracellular HIV-p24 staining and by HIV-p24 ELISA in cell-culture supernatant, as previously described (18, 27).

### Flow cytometry staining and Fluorescence activated cell sorting (FACS)

The following fluorochrome-conjugated Abs were used for flow cytometry analysis: HIV-p24 FITC (KC57) (Beckman Coulter, Fullerton, CA, USA), HIV-p24 PE (KC57) (Beckman Coulter, Fullerton, CA, USA), CD3 Pacific blue (UCHT1), CD4 PerCP/Cy5.5 (RPA-T4) (Biolegend, San Diego, CA, USA), CD4 Alexa Fluor 700 (RPA-T4), CCR6 PE (11A9), CD45RA Alexa eFluor 780 (HI100), RORC2 Alexa Fluor 647 (Q31-378), Ki-67 BUV395 (B56), IL-17A PE (eBio64DEC17) and IFN-γ Alexa Fluor 700 (B27). Live/Dead Fixable Aqua Dead Cell Stain Kit (Vivid, Life Technologies, Burlington, Ontario, CA) was used to exclude dead cells. Intracellular staining was performed using the BD Cytofix/Cytoperm kit (BD Biosciences) and intranuclear staining was performed using the eBioscience Foxp3/Transcription Factor Staining Buffer Set. Cells were analysed with the BD-LSRII cytometer, BD LSRFortessa and BD-Diva (BD Biosciences) and FlowJo version 10 (Tree Star, Inc., Ashland Oregon, USA). The positivity gates were placed using fluorescence minus one (FMO) strategy (8, 20). For FACS, memory CD4 T cells from PBMCs of ART+ PLWH were isolated by negative selection using magnetic beads. CCR6^+^RORC2+, CCR6+RORC2− and CCR6^−^RORC− T cells were sorted by FACS (BDAria II; BD Biosciences) using the antibodies CD3 Pacific blue (UCHT1), CCR6 PE (11A9), CD45RA Alexa eFluor 780 (HI100), and RORC2 Alexa Fluor 647 (Q31-378).

### ELISA

HIV-p24 levels in cell culture supernatant were quantified using a homemade sandwich ELISA, as described previously (8, 18). Briefly, virions in cell supernatants were lysed using a homemade buffer solution (PBS 1X, Tween 20 0.05%, Triton X-100 2.5%, Trypan Blue 1% and Thimerosal 0.02% in deionized water) for 1h at 37°C. Levels of IL-17A and IFN-γ were measured in the cell culture supernatant, according to the manufacturer’s protocols (Thermofisher).

### Compounds

GSK2793955A, GSK2805956A, GSK2833332A, GSK2837269A and GSK2837270A were provided by GlaxoSmithKline at 100 mM stock in DMSO or freeze-dried and reconstituted in DMSO. GSK2691805A was synthesized in house according to (33) (see also Supporting Materials and Methods).

### Homogenous Time Resolved Fluorescence (HTRF) RORC2 Ligand Binding Assay

The Homogenous Time Resolved Fluorescence (HTRF) RORC2 Ligand Binding assay measures the interaction of co-factor SRC1 peptide with a purified bacterial-expressed RORγ ligand-binding domain (LBD). This assay is based on the knowledge that nuclear hormone receptors interact with cofactors in a ligand dependent manner. RORγ has a basal level of interaction with the co-activator SRC1 in the absence of ligand, thus it is possible to find ligands that enhance or inhibit the RORγ/SRC1 peptide interaction. The sites of interaction have been mapped to LXXLL Leucine Charge Domain 2 motifs in the co-factor sequence, and to the AF2 domain of the nuclear receptor. Short peptide sequences containing the LXXLL motif mimic the behaviour of full-length co-factors. A biotinylated SRC1 peptide residues 676-700 (CPSSHSSLTERHKILHRLLQEGSPS-CONH2) was used as an inhibitor (i.e. ‘cold peptide’) to compete-off the biotinylated peptide from the RORγ LBD. The biotinylated peptide can be displaced by the unbiotinylated form. For this assay, an equal volume of biotinylated SRC1 peptide/Europium-labeled streptavidin (Perkin Elmer) was added to biotynlated RORγ/APC-labeled strepavidin (Perkin Elmer), each in 10 mM DL-dithiothreotol (DTT, JT Baker) with 400 nM D-biotin (Pierce), to give a final solution of 20 nM biotinylated-SRC1 peptide, 1 nM Europium-streptavidin, 20 nM biotinylated-RORγ, 10 nM APC-streptavidin in 10 mM DTT with 400 nM biotin. After a 5 minute incubation at room temperature, 25 μL of the peptide/RORγ solution was added to 384-well assay plates containing 1 μL of titrations of compounds in 100% DMSO. Plates were incubated for one hour and read on ViewLux ultra HTS Microplate Imager (Perkin Elmer) in Lance mode for EU/APC. For dose response curves the APC counts (@671) were divided by the europium (@618) counts to compensate for quenching effects and account for well to well variation due to liquid handling errors. Data were normalized using the following equation: (unknown- ave basal)/ (ave basal - ave background) * 100 = % activation or inhibition. A response of 0% would be inactive, >0% indicates enhancement of the protein peptide interaction and < 0% (negative values) indicates inhibition of the protein peptide interaction. Results were analyzed with ActivityBase (IDBS) using a 4-parameter fit equation.

### Jurkat RORC2 IL-17F Promoter Luciferase Reporter Assay

The Jurkat RORC2 IL-17F Promoter Luciferase Reporter Assay measures RORC2-specific (human IL-17 conserved non-coding sequence (CNS)) promoter driven luciferase-reporter activity to indirectly assess RORC2 activity. Both RORC gene and reporter construct were sequentially transfected into Jurkat cells and stably integrated. The Jurkat double stable cell line expressing the human RORC2 and IL-17 luciferase-reporter construct was adjusted to a concentration of 0.5 x 10^6^ cells/mL in medium (RPMI-1640, 10% FBS, 2 mM Glutamax) and mixed with 166 ng/mL of anti-CD3 antibody (GlaxoSmithKline). 50 μL of cells/antibody was added to each well of 384-well plates containing compound titrations in 100% DMSO and plates were incubated at 37 °C for 18 h. 20 μL of Steady-Glo Luciferase Assay reagent (Promega) was added to each well and plates were incubated for 30 minutes at room temperature. Luminescence was measured on a ViewLux ultra HTS Microplate Imager (Perkin Elmer) and data were analyzed with ActivityBase (IDBS) fitting a 4-parameter equation.

### Cell lines and viruses

293T cells (European Collection of Authenticated Cell Culture [ECACC] Public Health England, UK) were grown in Dulbecco’s modified Eagle’s medium (DMEM) (Gibco Labs, Paisley, UK) supplemented with 10% fetal calf serum (FCS) (Helena Bioscience, Newcastle, UK) and 2 mM glutamine (Gibco Labs) at 37°C in 5% CO_2_. Jurkat E6.1 (ECACC) and Jurkat indicator line 1G5 containing the firefly luciferase gene driven by the HIV LTR (AIDS Research and Reference Reagent Program, Division of AIDS, NIAID, NIH: from Dr. Estuardo Aguilar-Cordova and Dr. John Belmont) were grown in RPMI medium (Gibco Labs) supplemented with 10% FCS at 37°C in 10% CO_2_. Jurkat cell line IL-17CNS luciferase clone 539 (GSK BIOCAT128253) (here called Jurkat Luc) were generated by transfection of Jurkat E6.1 cells with plasmid pGL4-huIL-17 3-K CNS promoter containing the luciferase gene driven by the 1 Kb IL-17 CNS-5 enhancer fused to the 2 Kb promoter regions and were grown in RPMI medium with 10% FCS and 200μg/ml hygromycin at 37°C in 10% CO_2_. Viral stocks were prepared by Fugene transfection of 293T cells as described previously (38) using pHIV LAI∆env (gift of Michael Emerman, Fred Hutchinson Cancer Research Centre, Seattle, USA) and pMD.G expressing VSV-G or using HIV isolate NL4.3 (Centre for AIDS Reagents, Health protection Agency, UK). Supernatant containing viral particles was collected 48h and 72h post-transfection as described (38, 64). For infections, 13ml of 1G5 indicator cells (~0.9×10^6^/ml) were mixed with 2ml NL4-3 supernatant. The mix was dispensed robotically, 45 μl/well onto 384-plates preloaded with drug dilutions. Samples were analysed 48 h post-infection using the BrightGlo assay according to the manufacturer’s instructions in a Pherastar plate reader. To generate Jurkat and 293T cells stably expressing human RORC2, the myc-DDK-tagged-RORC2 cDNA was PCR-amplified from plasmid RC212239 (Origene) and cloned into the MLV-based retroviral vector pMIG Blasti (gift of Jeremy Luban, UMASS, USA). Virus was produced in 293T cells as described and used to transduce Jurkat or 293T cells, which were selected in the presence of 5 μM blasticidin for 10 days.

### Jurkat KD RORC2

To generate 1G5 Jurkat indicator cells with a stable KD of RORC2, four shRNA hairpins were obtained from Sigma mission catalogue 3-1245h1C1; 4-1036h21C1; 4-363h21C1 and 3-978h1C1 and were cloned into lentiviral vector pLKO.1-puro (Addgene, catalogue #8543). Lentiviral vectors were produced in 293T cells by Fugene transfection and the supernatant was used to infect Jurkat cells. Forty-eight hours post-infection, cells were selected with 5 μM puromycin for 4 days.

### Western blot

2×10^6^ cells were centrifugated (100g, 5minutes), washed with ice cold phosphate buffered saline (PBS) once and lysed with 150μl of SDS sample buffer (140mM Tris pH8, 2% SDS, 50 mM DTT, 0.5M sucrose, 2mM MgCl_2_, Bromophenol-blue). Lysates were incubated at 100°C for 5 minutes. Samples were loaded to a NuPAGE™ Bis-Tris Protein Gel. Western blotting was performed following wet electrophoretic transfer (1h, 100mV) to a PVDF membrane. After probing with the primary antibody at 4°C overnight, HRP conjugated secondary antibodies were used for detection by luminescence. Primary antibodies were: rabbit polyclonal anti-RORC/RORC2 cat. LS-C40832, rabbit polyclonal anti-RORC/RORC2 L-C116877 both from LSBio (Seattle, WA) and mouse mAb anti-c-Myc (ThermoFisher, 9E10).

### qPCR

For Taqman qPCR, approximately 1×10^6^ Jurkat cells were washed twice in PBS and total DNA was extracted with the Qiamp® DNA Minikit (Qiagen, Manchester, UK). Quantitative PCR reactions were carried out as previously described using 0.3pmol each primer and 0.15 pmol of the probe in 25 μL volume containing 100-300ng total DNA using an ABI Prism^®^ 7000 Sequence Detection System (SDS). For amplification of (−) DNA strand (GFP), primers used were forward: CAACAGCCACAACGTCTATATCAT, reverse ATGTTGTGGCGG ATCTTGAAG and probe 5’-FAM-CCG ACA AGC AGA AGA ACG GCA TCA A-3’TAMRA. For amplification of 2LTR circular DNA, the same conditions were used with primers 2LTRqPCR-F: 5’-AACTAGAGATCCCTCAGACCCTTTT-3’ and 2LTRqPCR-RC: 5’- CTTGTCTTCGTTGGGAGTGAATT-3’ and 2LTR probe 5’-FAM-CTAGAGTTTTCCACACTGAC-0-TAMRA-3’. Standards were prepared by PCR amplification of DNA from acutely infected cells with primers 2LTRF 5’-GCCTCAATAAAGCTTGCCTGG-3’ and 2LTRRC 5’-TCCCAGGCTCAGATCTGGTCTAAC-3’. The amplification product was cloned into TOPO vector, amplified and confirmed by sequencing (64). Alu-LTR Taqman qPCR was carried out as previously described (64) using primers ALU-forward, AAC TAG GGA ACC CAC TGC TTA AG and LTR1-reverse, TGC TGG GAT TAC AGG CGT GAG (for first round amplification) and ALU-forward AAC TAG GGA ACC CAC TGC TTA AG, LTR2-reverse, TGC TAG AGA TTT TCC ACA CTG ACT, ALU-probe, FAMRA – TAG TGT GTG CCC GTC TGT TGT GTG AC – TAM (for second round Taqman qPCR). Integrated HIV-DNA in human primary memory CD4^+^ T-cells was quantified by nested real-time PCR using specific primers and amplification conditions as previously described (8, 20).

### RT-qPCR

Jurkat cells were grown in RPMI + 10% FCS. 1.5 x 10^6^ cells/ml were seeded onto 12-well plates. Cells were infected with HIV-1 LAI_GFP_ and 24 hours cells an aliquot was analyzed by FACS and another aliquot was treated with GSK2837269A (5 μM) or DMSO. After 24 hours, cells were collected, and RNA was isolated using RNAesy mini kit (Qiagen) and reverse transcribed using SuperScript™ III Reverse Transcriptase (Invitrogen) after digestion with RNAse-free DNAse I (Promega). Quantitative TaqMan qPCR was performed using GFP forward primer (5’-CAA CAG CCA CAA CGT CTA TAT CAT-3’), GFP reverse primer (5’-ATG TTG TGG CGG ATC TTG AAG-3’) and GFP probe (5’-FAM-CCG ACAAGC AGA AGA ACG GCA TCA A- TAM-3’) in an Eppendorf MasterCycler Realplex. HIV-GFP mRNA expression was normalized to that of hu28S rRNA expression using hu28S rRNA forward primer (5’-TTG AAA ATC CGG GGG AGA G-3’) and hu28S rRNA reverse primer (5’-ACA TTG TTC CAA CAT GCC AG-3’). For GFP mRNA, the TaqMan PCR assay was set in a final volume of 20μl containing 250ng cDNA, 0.5μM of each primer, 0.15μM of GFP probe and TaqMan universal master mix II (2X) (Qiagen). For hu28S rRNA, the SYBR Green qPCR reaction was carried out in a final volume of 20μl using 250 ng cDNA, 0.4 μM hu28S rRNA forward and reverse primers and PowerUp™ SYBRTM Green Master Mix. For primary memory CD4^+^ T-cells, Jurkat, ACH2, TZM-bl and HT-29 cell lines shown in **Supporting information Figure 1**, total RNA was extracted using the RNeasy kit (Qiagen) and quantified by Pearl nanophotometer (Implen, Germany). RORC1 and RORC2 gene expression was evaluated by One step SYBR green real-rime RT-PCR (Qiagen) using a Light-Cycler 480 II as follows; reverse transcription at 50°C for 30 min, 15 min at 95°C and then 45 cycles each at 94°C for 10 s, 61°C for 10 s, and 72°C for 10 s. The sequence of primers used for RORC isoforms were: RORC2 rRNA forward primer, 5’--CTGCTGAGAAGGACAGGGAG-3’; RORC1 rRNA forward primer, 5’- CACAGAGACAGCACCGAGC-3’; RORC2/RORC1 rRNA reverse primer (same for both isoforms) 5’-AGTTCTGCTGACGGGTGC-3’. The relative expression of RORC was normalized relative to the internal control 28S. The sequence of primers used for 28S were 28s rRNA forward primer 5′-CGAGATTCCTGTCCCCACTA-3′ and 28s rRNA reverse primer, 5′ GGGGCCACCTCCTTATTCTA-3′. Primers were obtained from IDT. Each reaction was performed in triplicates. After real time amplification, melting curve analysis was used to determine the uniformity of the thermal dissociation profile for each amplification product.

### RNA interference in primary CD4+ T cells

RNA interference was performed as previously described by our group (65). Briefly, PBMCs were thawed and rested overnight at 37 °C. Memory CD4+ T-cells were isolated by negative selection using magnetic beads (Miltenyi Biotec), as described previously (8, 20). Cells were stimulated by CD3/CD28 Abs for 2 days and nuclofected with 100 μM RORC or non-targeting (NT1) siRNA (ON-TARGETplus SMART pool, Dharmacon) using the Amaxa Human T cell Nucleofector Kit (Amaxa, Lonza), according to the manufacturer’s protocol. Cells were suspended in the NF solution (100 μl/2×106 cells) and nucleofected using the Amaxa Nucleofector II Device and the human activated T-cell protocol (T-20). Cells (2×106) were transferred into 48-well plates containing 1 ml of RPMI1640 (10 % FBS, 5 ng/ml IL-2, w/o antibiotics) and cultured for another 24 hours at 37 °C before HIV exposure.

### ChIP-qPCR

ChIP assays were performed as described previously (64) with some modifications. Briefly, 5 × 10^7^ Jurkat cells expressing myc-tagged RORC2 or the pMI-blasti “empty” vector (EV) were infected with the VSV-G pseudotyped LAI∆env-GFP at an m.o.i. of 0.3. Twenty four hours later, cells were collected in 50-ml tubes and chemically cross-linked by the addition of 1/10th volume of fresh 11% formaldehyde solution added directly to cell culture media and incubated for 20 min at room temperature with gentle rotation followed by the addition of 1/20th volume of cold 2.5 M glycine and incubated at 4 °C for 5 min. Cells were then collected by centrifugation at 4 °C, and the pellet was rinsed twice with PBS and flash-frozen in liquid N_2_. Cells were resuspended and lysed in 1 mL lysis buffer 1 (50 mM Hepes-KOH, pH 7.5, 140 mM NaCl, 1 mM EDTA, 10% glycerol, 0.5% IGEPAL, 0.25% Triton X-100) for 10 min at 4 °C with slow rotation. Nuclei were pelleted by centrifugation and gently resuspended in 1 mL Nuclei Wash buffer (200 mM NaCl, 1 mM EDTA) for 10 mins at 4 °C with gentle rotation. Nuclei were pelleted by centrifugation and 1 mL of Lysis Buffer 3 (LB3) (10 mM Tris, pH 8, 100 mM NaCl, 1 mM EDTA, 0.5 mM EGTA, 0.1% sodium deoxycholate, 0.5% N-laurylsarcosine) was added without disturbing the pellet followed by incubation for 10 mins at 4 °C with gentle rotation. This step was repeated once then nuclei were pelleted by centrifugation, resuspended in 300 μL LB3 and kept on ice for 10 mins. Samples were sonicated using a Diagenode Bioruptor device (30s pulse/90 sec pause x 10 cycles). Samples were centrifuged at 13,000 rpm for 6 mins at 4 °C and supernatant was collected in pre-cooled 1.5 mL tubes. The resulting whole-cell extract (WCE) was incubated overnight at 4 °C with 100 μL of protein-G magnetic Dynabeads preincubated with 10 μg of the appropriate antibody for 3 h on rotating platform (9 rpm) in a cold room. Antibodies used were: rabbit polyclonal anti-H3K9me3 (abcam, ab8898), normal rabbit IgG (Merk Millipore, 12-370), mouse mAb anti-c-Myc (ThermoFisher, 9E10) and rabbit polyclonal anti-NF-κB p65 acetyl K310 (Abcam ab19870). The next day beads were washed (5 mins with slow rotation) two times with low salt buffer (10mM Tris HCl pH 8, 150 mM NaCl, 1mM EDTA, 1% Triton ×100, 0.1 % SDS, PMSF) then once with high salt buffer (10mM Tris HCl pH 8, 500 mM NaCl, 1mM EDTA, 1% Triton ×100, 0.1 % SDS, PMSF) then once with LiCl buffer (10mM Tris HCl pH 8, 1mM EDTA, 0.5 mM EGTA, 250 mM LiCl, 1% IGEPAL, 1% NaDOC, PMSF) then once with TE buffer and finally with elution buffer (TE + 1% SDS). Bound complexes were eluted from the beads by heating at 65 °C with occasional vortexing and cross-linking in the immunoprecipitation and WCE samples was reversed by incubating at 65 °C for 6–7 h. Immunoprecipitation and WCE DNA was then purified by treatment with RNase A, proteinase K and extracted with phenol/chloroform/isoamyl alcohol extractions. ChIP products were quantified by Real Time qPCR. Primer sequences were as follows: IL-17 enhancer forward: 5’ TGATAGCCCAACCACAATGTG – 3’ (IL-17 gene nt 1051 - 1072). IL-17 enhancer reverse: 5’- ACCTATACGTTAGCAGGCACA – 3’ (IL-17 gene nt 1220 - 1241). IL-17 promoter forward: 5’- TCTGCCCTTCCCATTTTCCT-3’ (IL-17 gene nt 2886 – 2906) IL-17 promoter reverse: 5’- ATGGATGAGTTTGTGCCTGC-3’ nt 3064 – 3084. Nuclear hormone receptor responsive element (NRRE-1) in the HIV-1 LTR Forward 5’- TCTACCACACACAAGGCTACT-3’, Reverse 5’-ACAAGCTGGTGTTCTCTCCT-3’; RORC2 consensus sequence (CS) in HIV-1 Pol forward: 5’-GGGAAAGCTAGGGGATGGTT-3’ (HIV-1 nt 5137 - 5157), RORC2 CS reverse: 5’-TCAGGGTCTACTTGTGTGCT -3 (HIV-1 nt 5322-5342). Real time PCR was carried out in an Eppendorf Realplex in a final volume of 20 μL containing 1x SYBR green master mix (Applied Biosystems), 0.4 μM each primer and 2 μL DNA (pre-diluted 1:10). Cycles parameters were: 95 °C for 2 minutes for 1 cycle, followed by 95 °C for 1 minute, 55 °C for 55 seconds and 65 °C for 1min 30 seconds for 45 cycles. ChIP signal was calculated using the percent input method.

### Viral outgrowth assay

A simplified VOA was performed, as we previously described (50). Briefly, memory CD4+ T-cells were cultured at 1×10^6^ cells/well in 1 ml of media (RPMI, 10% FBS, 1% Penicillin/Streptomycin) in a 48-well plate (Costar) coated with CD3 Abs (1 μg/ml; BD Biosciences, Clone UCHT1) and in the presence of soluble CD28 Abs (1 μg/ml; BD Biosciences, Clone CD28.2). At day 3 post-stimulation, cells from each original replicate were individually washed with media, split into two new CD3 Abs-uncoated 48-well plate wells, and cultured in media containing IL-2 (5 ng/ml; R&D Systems), in the presence or in the absence of GGSK2691805A (5 μM). Cells from each well were further split into two new wells (without washing) at day 6 and 9 post-stimulation, with 50% of the media being refreshed with IL-2 with/without GSK2691805A. Cells were kept in culture for a total of 12 days.

### Statistical Analysis

Statistical analyses were performed with GraphPad Prism 7 software (GraphPad Software, Inc., La Joya, CA, USA). One-way ANOVA and Friedman along with Dunnett’s and Dunn’s multiple comparisons test respectively evaluated the statistical differences. The use of non-parametric tests is justified by the fact that data sets did not pass the normal distribution test Kolmogorov-Smirnov (data not shown). P-values of ≤0.05 were arbitrarily considered statistically significant.

## Acknowledgments

This work was supported by a grant from Neomed/GSK (to AF and PA), the EU HEALTH-F3-2012-305137 HIVINNOV grant (to AF), the Canadian HIV Cure Enterprise Team Grant (CanCURE 1.0) funded by the Canadian Institutes of Health (CIHR) in partnership with CANFAR and IAS (CanCURE 1.0; # HIG-133050 to PA); The Canadian HIV Cure Enterprise Team Grant (CanCURE 2.0) funded by the CIHR CIHR (#HB2-164064, and PJT #153052 to PA and and CIHR # MOP 103230 and PTJ 166049 to JPR). TRWS and DP received doctoral fellowships from the Université de Montréal and Fonds de Recherche en Santé Québec (FRQ-S). JPR holds the Louis Lowenstein Chair in Hematology and Oncology, McGill University.

We thank Veronique Birault and David Favre for their support and helpful comments, David Irlbeck for support with characterization of the compounds, David Selwood for advice on medicinal chemistry and Robin Ketteler for access to the MRC Translational Resource Centre facilities. The authors thank Dr. Dominique Gauchat and Philippe St Onge (Flow Cytometry Core Facility, CHUM-Research Center, Montréal, QC, Canada) for expert technical support with polychromatic flow cytometry sorting; Olfa Debbeche (Biosafety Level 3 Core Facility CHUM-Research Center, Montréal, QC, Canada); Mario Legault (FRQ-S/AIDS and Infectious Diseases Network; Montréal, QC, Canada) for help with ethical approvals and informed consents, and Josée Girouard, Angie Massicotte, and Maria Fraraccio (McGill University Health Centre, Montréal, QC, Canada), for their key contribution to blood collection from HIV-infected study participants and clinical information from HIV-infected and uninfected donors. The authors acknowledge the key contribution of all study participants for their precious gift of leukapheresis essential for this study.

## Authors’ contributions

TRWS and YZ designed and performed research, analysed data, prepared figures, and contributed to writing the manuscript. DS, AZ, LRM, DP, ML, DC, KK, and SO designed and performed experiments, analysed data and prepared figures. JPR provided access to clinical samples/information, set up clinical research protocols, and contributed to manuscript revision. HA-M provided critical reagents and expertise. PA and AF conceived the research study, designed research, analysed data, and wrote the manuscript. All co-authors revised and approved the manuscript.

## Materials availability

GSK compounds are available upon reasonable request. GSK2691805 is commercially available.

