## Supplemental information for "Th17 cell master transcription factor RORC2 regulates HIV-1 gene expression and viral outgrowth"

**Supporting information Table 2: Clinical parameters of HIV-infected untreated and ART-treated study participants.**

| ID | Sex | Age″ | CD4 count# | CD4:CD8 Ratios | Viral load& | Time since infection* | ART | Time on ART* |
|---|---|---|---|---|---|---|---|---|
| ART+ #1 | M | 45 | 318 | 0,7 | <40 | 150 | Delaviridine Kivexa | 50 |
| ART+ #2 | M | 44 | 459 | 0,8 | <40 | 189 | Truvada Raltegravir | - |
| ART+ #3 | M | 46 | 581 | 0,5 | <40 | 99 | - | 94 |
| ART+ #4 | M | 32 | 523 | 1,0 | <40 | 52 | Truvada Reyataz | 48 |
| ART+ #5 | M | 57 | 514 | 0,9 | <40 | 16 | Tivicay Truvada | 11 |
| ART+ #6 | M | 44 | 398 | 0,5 | <40 | 154 | Complera | 25 |
| ART+ #7 | M | 36 | 542 | 0,7 | <40 | 13 | Stribild | 12 |
| ART+ #8 | M | 49 | 458 | 0,5 | <40 | 227 | Truvada Viramune | 201 |
| ART+ #9 | M | 30 | 598 | 1,0 | <40 | 80 | Stribild | 77 |
| ART- #1 | M | 24 | 316 | 0,5 | 9,496 | 55 | None | N.A |
| ART- #2 | M | 47 | 529 | 1,2 | 3,189 | 110 | None | N.A |
| ART- #3 | M | 42 | 221 | 0,5 | 41,774 | 89 | None | N.A |
| ART- #4 | M | 50 | 389 | 0,3 | 97,552 | 150 | None | N.A |
| ART- #5 | M | 40 | 1,068 | 1,2 | 22,812 | 2 | None | N.A |
| ART- #6 | M | 24 | 897 | 1,3 | 18,621 | - | None | N.A |

M, male; F, female; ART-, ART-untreated PLWH; ART+, ART-treated PLWH; ″, years; #, cells/µl; &, HIV-RNA copies/ml plasma; *, months on ART; -, information not available; N.A, not applicable

### Supporting information Figures

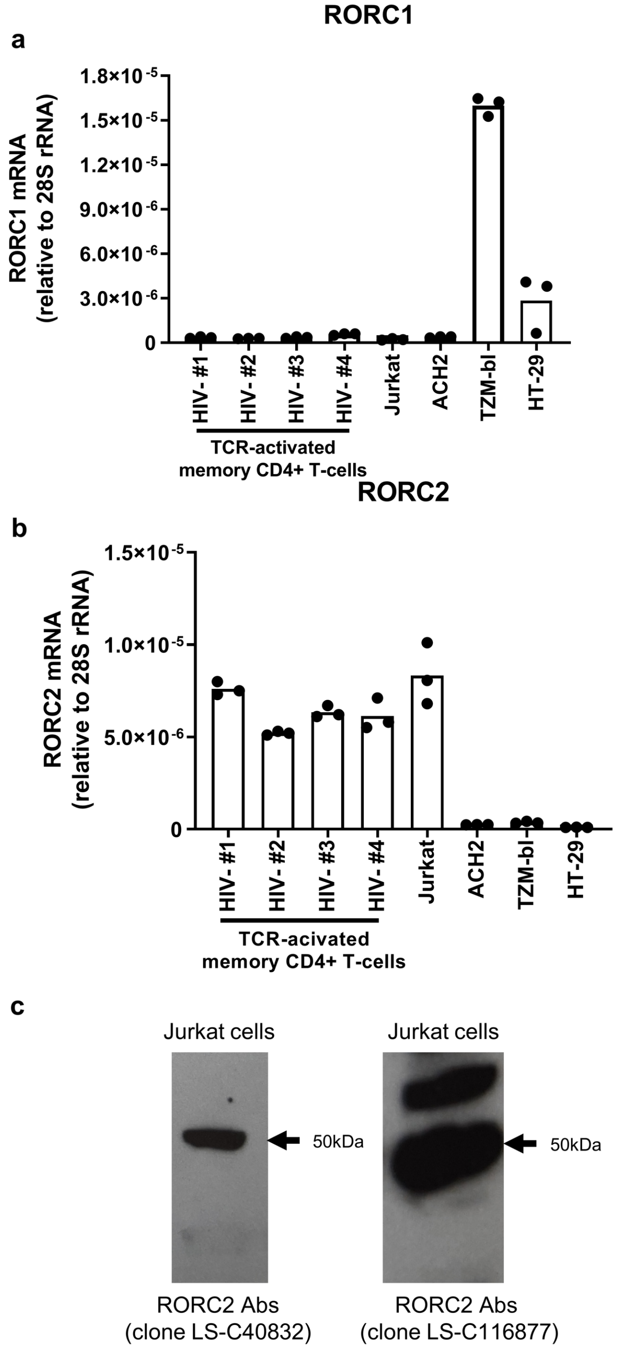


**Supporting information Figure 1. RORC2, but not RORC1, is expressed in primary CD4+ T-cells and Jurkat cells. (a-b)** The relative gene expression of RORC1 mRNA **(a)** and RORC2 mRNA **(b)** was evaluated by real-time RT-qPCR in CD3/CD28-activated memory $CD4^+$ T-cells isolated from n=4 HIV-uninfected individuals as well as the Jurkat, ACH2, TZM-bl and HT-29 cell lines. **(c)** Western blot of Jurkat cell lysates to detect RORC2 expression. The antibody used is indicated below each panel.

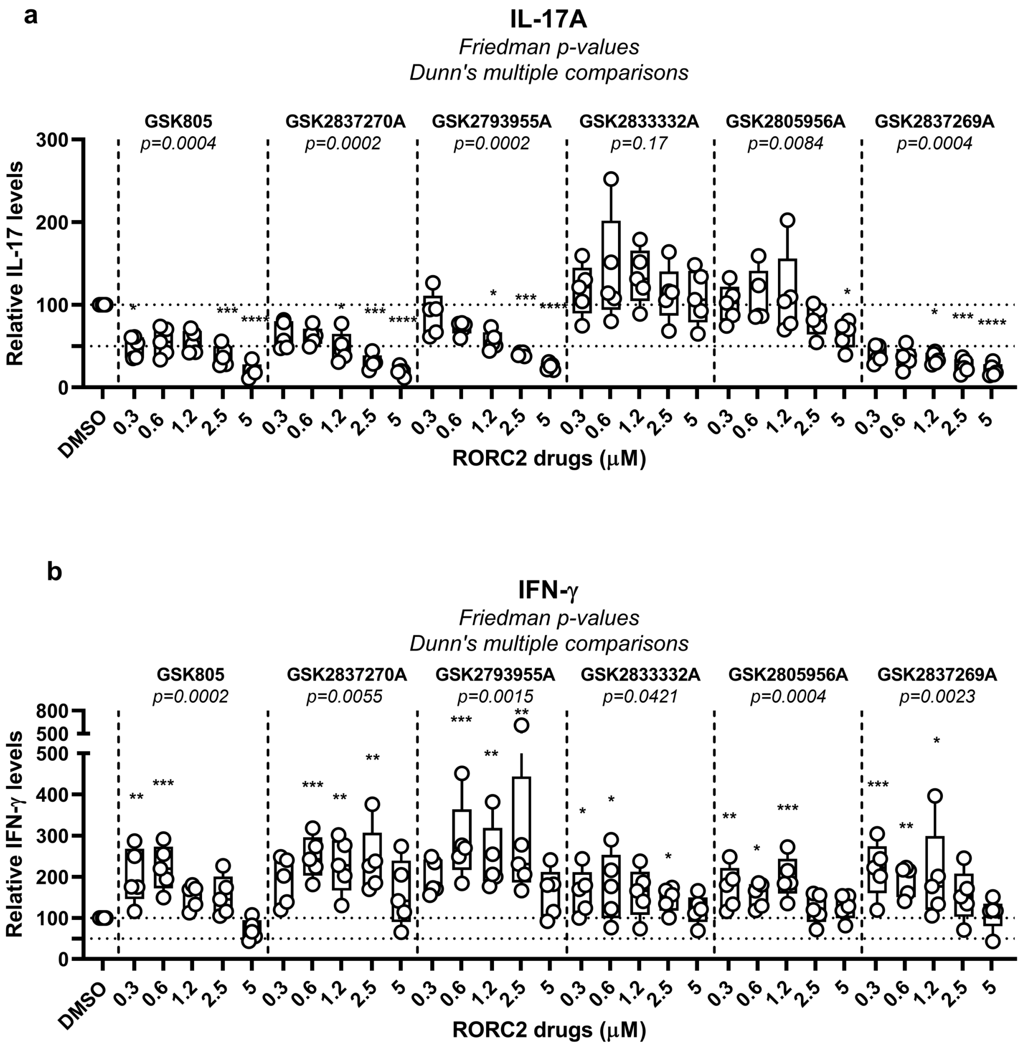


**Supporting information Figure 2. RORC2 inhibitors reduce IL-17A, but not IFN-γ production.** Memory CD4+ T-cells were isolated and stimulated with CD3/CD28 Abs in the presence or absence of the indicated RORC2 inhibitors, as described in Figure 2. Levels of IL-17A and IFN-γ were measured by ELISA at day 3 post-stimulation. Statistical analysis of IL-17A **(a)** and IFN-γ **(b)** levels in cell culture supernatants of experiments performed with cells from n=5 HIV-uninfected individuals. Friedman p-values, with Dunn's multiple comparison significance, are indicated on the graphs (*, $p<0.05$; **, $p<0.01$; ***, $p<0.001$; ****, $p<0.0001$).

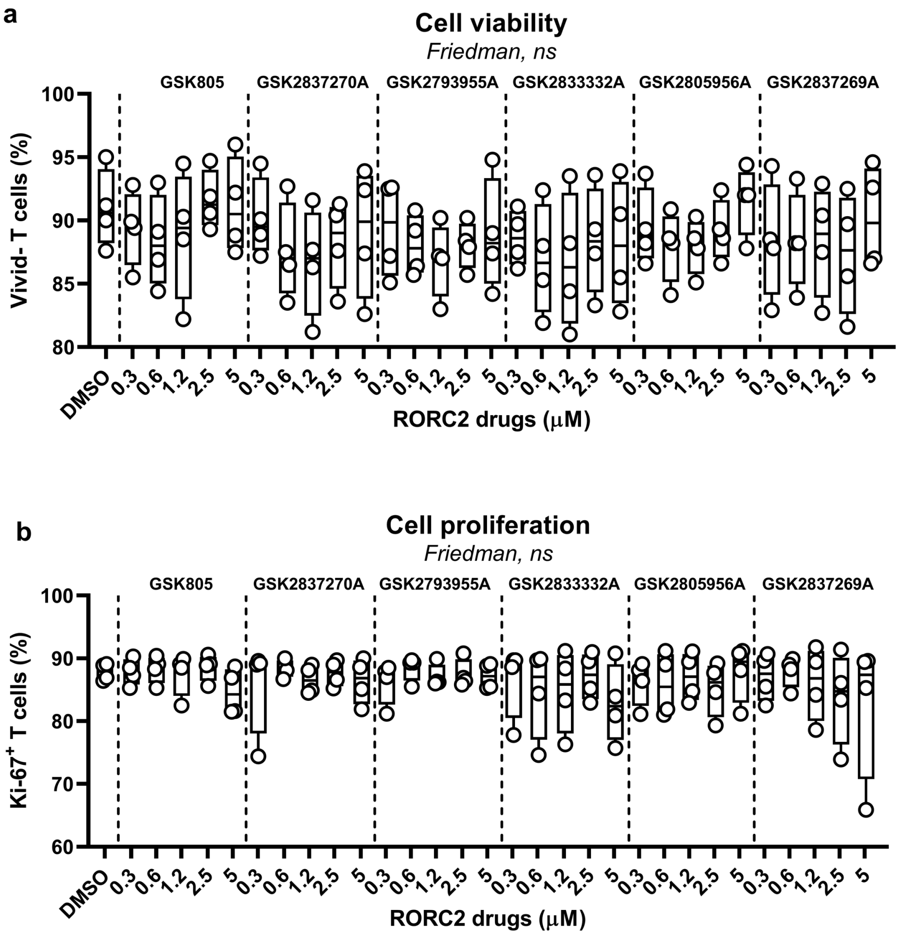


**Supporting information Figure 3. No effect of RORC2 inhibitors on cell viability and proliferation.** A fraction of memory $CD4^+$ T cells from experiments depicted in Figure 2 were harvested 3 days after CD3/CD28 stimulation and stained with Live/dead Fixable Aqua dead dye and fluorochrome-conjugated Ki-67 Abs for flow cytometry analysis. Shown are the statistical analysis of the frequency of Live (Vivid-) **(a)** and Ki-67+ cells **(b)** in experiments performed with cells from n=5 HIV-uninfected individuals.

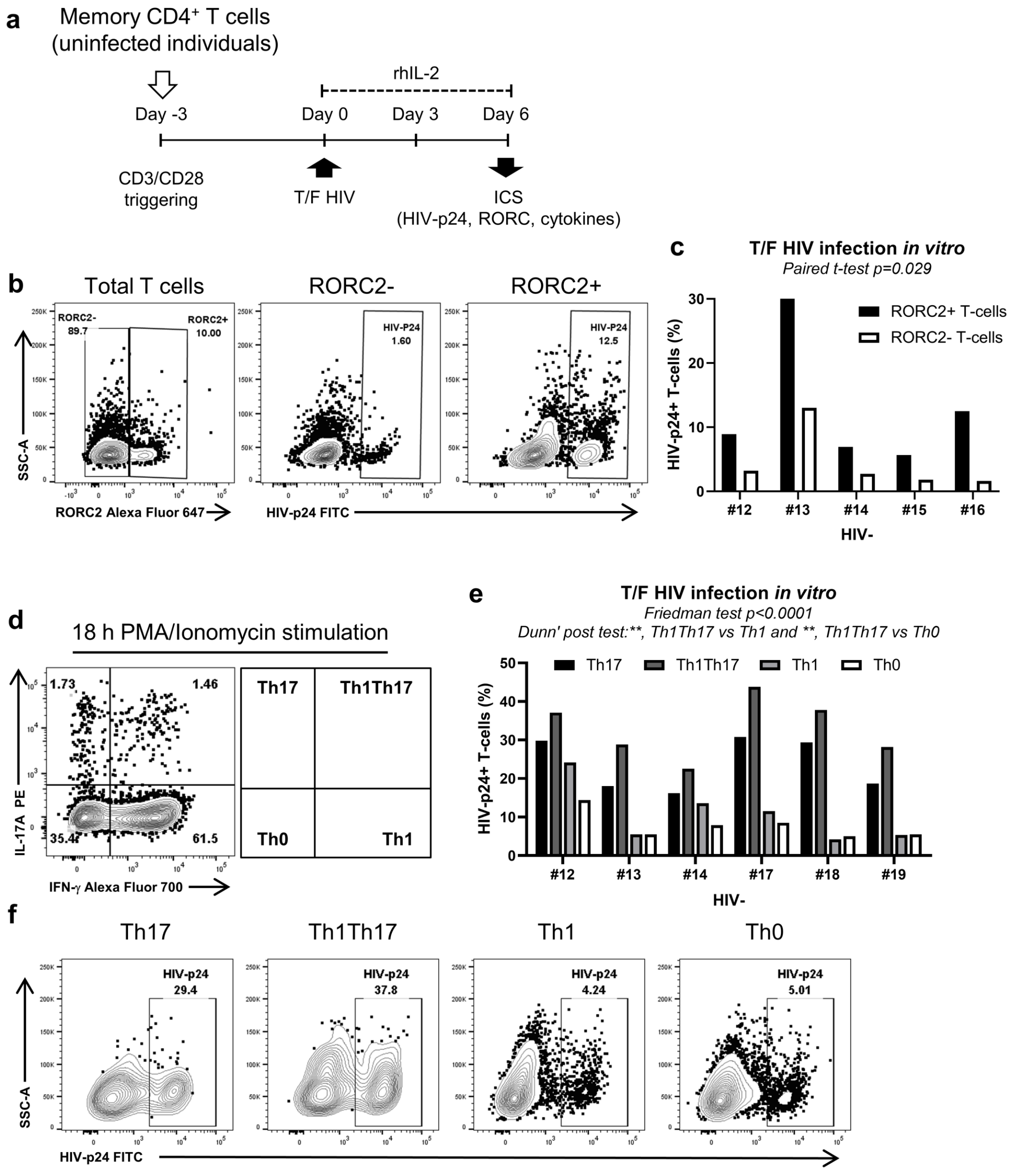


**Supporting information Figure 4**. **Preferential HIV-1 replication in Th17-polarized RORC2+ cells *in vitro*. (a).** Experimental flow chart; memory $CD4^+$ T-cells isolated from n=5 HIV-uninfected individuals were stimulated with CD3/CD28 Abs for 3 days and exposed to HIV$_{THRO}$. After infection, cells were cultured in the presence of IL-2 for 6 days. The intracellular expression of HIV-p24, RORC2 and cytokines (IL-17A and IFN-γ) was analysed by flow cytometry. Shown are contour plots of RORC2 expression **(b, left panel)** and HIV-p24 expression in RORC2+ and RORC2- cells of one representative individual **(b, middle/right panels)** and statistical analysis of results obtained with cells form n=5 individuals **(c)**. **(d)** Gating strategy used to identify Th17 (IL-17A+IFN-γ-), Th1Th17 (IL-17A+IFN-γ+), Th1 (IL-17A-IFN-γ+), Th0 (IL-17A-IFN-γ-)**. (e)** Contour plots of HIV-p24 expression in Th subsets of one representative individual and **(f)** statistical analysis of the frequency of HIV-p24 cells in Th subsets for experiments performed with cells from n=6 individuals.

```
IIIB            CAGGAGAAAGAGACTGGCATTTGGGTCAGGGAGTCTCCATAGAATGGAGGAAAAAGAGAT 4866
NL4-3           CAGGAGAAAGAGACTGGCATTTGGGTCAGGGAGTCTCCATAGAATGGAGGAAAAAGAGAT 5320
C.96BW06.H51    CAGGAGAAAGAGAGTGGCATTTGGGTCATGGAGTCTCCATAGAATGGAGATTGAGAAAAT 5310
RORC            ------------------AWNTAGGTCA-------------------------------- 10
                                  **********
```

RORC2 binding motif

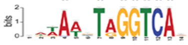

**Supporting information Figure 5**. RORC2 DNA consensus sequence in HIV-1 Pol. The consensus sequence (Ciofani et al. 2012) was aligned using ClustalW2 to the proviral genome sequence of HIV-1 IIIB, NL4.3 and subtype C 96BW06.H51.

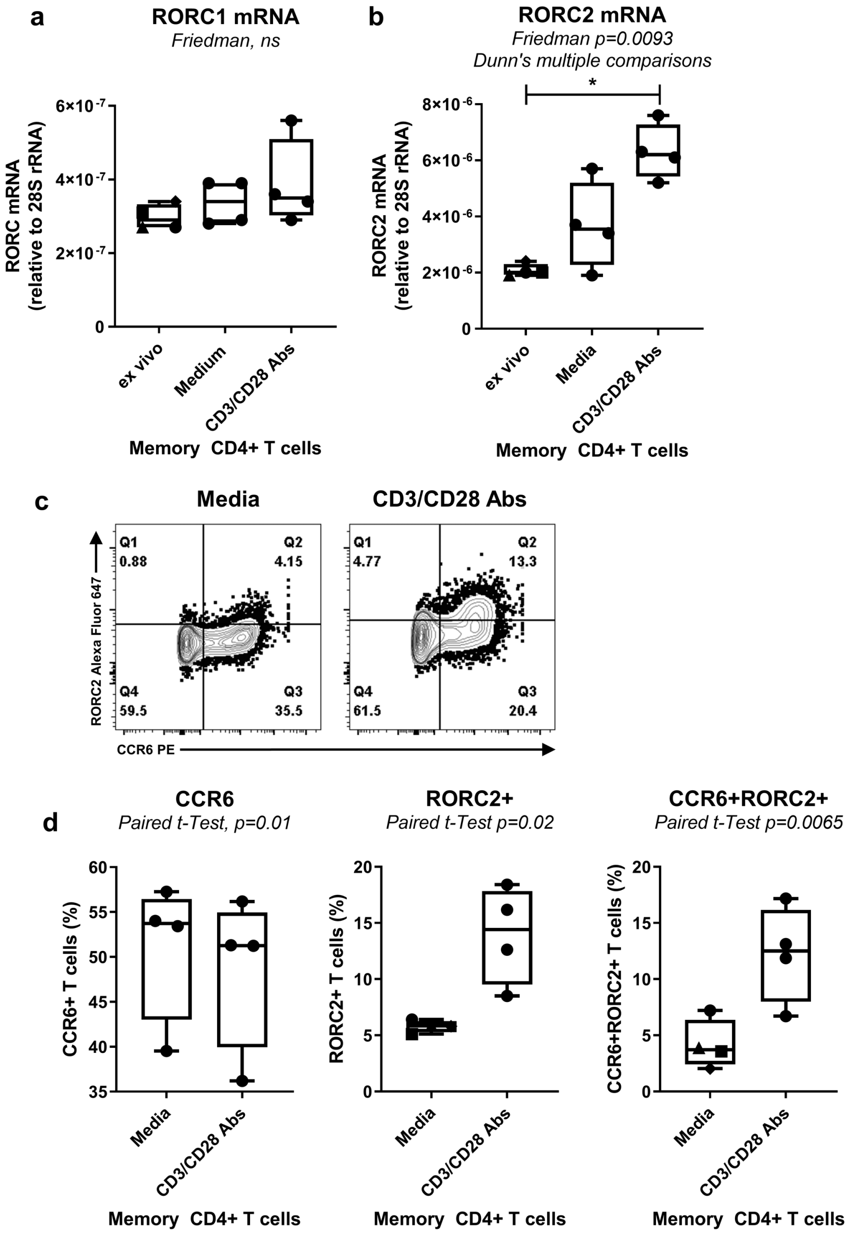


**Supporting information Figure 6. TCR triggering promotes RORC2 expression in primary CD4$^+$ T-cells without interfering with CCR6 expression.** Memory CD4$^+$ T-cells isolated from n=4 HIV-uninfected individuals were stimulated with CD3/CD28 antibodies for 5 hours to evaluate RORC1 and RORC2 mRNA expression by real-time RT-PCR and for 24 hours to evaluate RORC2 and CCR6 expression by flow cytometry. Shown are the statistical analysis of RORC1 **(a)** and RORC2 mRNA **(b)** expression *ex vivo*, and in CD3/CD28-activated and non-activated (medium) T cells. **(c)** Gating strategy and frequency of cells expressing RORC2 and/or CCR6 in one representative individual and **(d)** statistical analysis of the frequency of CCR6+, RORC2+ and CCR6+RORC2+ cells in n=4 individuals.

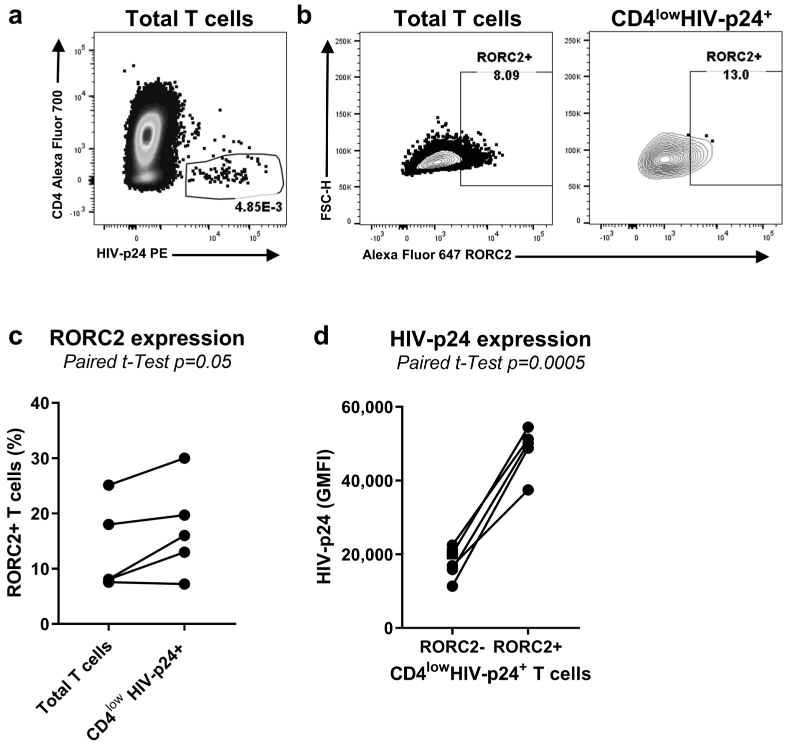


**Supporting information Figure 7. Higher RORC2 expression in memory CD4+ T-cells of ART-PLWH carrying translationally-competent HIV reservoirs.** Memory CD4+ T-cells isolated from n=5 ART- PLWH individuals were stimulated with CD3/CD28 antibodies for 3 days in the presence of antiretroviral drugs (raltegravir 0.2 μM and BMS806 1 μM) to prevent HIV cell-to-cell transmission *in vitro*. Intracellular expression of HIV-p24 and RORC2 was quantified by flow cytometry upon staining with fluorochrome-conjugated CD3, CD4, RORC2 and HIV-p24 antibodies. **(a)** Contour plots for CD4 and HIV-p24 expression, with $CD4^{low}HIV\text{-}p24^{+}$ T-cells identified as productively infected cells. **(b)** RORC2 expression in total T-cells (CD3+) and $CD4^{low}HIV\text{-}p24^{+}$ T-cells in one representative individual. **(c)** Statistical analysis of the frequency of RORC2+ T-cells n=5 individuals and **(d)** the geometric mean fluorescence intensity (GMFI) of HIV-p24 expression in RORC2- and RORC2+ within $CD4^{low}HIV\text{-}p24^{+}$ T-cells. Paired t-Test values are indicated on the graphs

**Supplementary Materials and Methods**

Synthesis of GSK GSK2691805A

$^1$H NMR spectral data were recorded on a Bruker 400 NMR spectrometer operating at 400 MHz. $CDCl_3$ is deuterochloroform, DMSO-d6 is hexadeuterodimethyl sulfoxide. Chemical shifts are given in parts per million ($\delta$) downfield from the NMR solvent. Abbreviations for NMR data are as follows: s = singlet, d = doublet, t = triplet, q = quartet, m = multiplet, dd = doublet of doublets, dt = doublet of triplets, app = apparent, br = broad. Step 1: To a solution of ethyl 2-(4-ethylsulfanylphenyl) acetate (1g, 4.7mmol) in dichloromethane (50mL) at 0°C was added 3-chloroperbenzoic acid (2.4g, 14.0mmol) and the reaction was stirred for 16 h at room temperature. The reaction was then quenched with saturated aqueous sodium carbonate solution (50mL) and extracted into dichloromethane (2 x 30mL). The organics were separated, dried ($MgSO_4$) and reduced *in vacuo*. Purification on the Biotage gave ethyl 2-(4-ethylsulfonylphenyl) acetate (980mg, 81% yield) as a gum. $^1$H NMR ($CDCl_3$): δH 1.25-1.30 (m, 6H), 3.11 (q, 2H), 3.71 (s, 2H), 4.18 (q, 2H), 7.49 (d, 2H) and 7.86 (d, 2H).

Step 2: To a solution of ethyl 2-(4-ethylsulfonylphenyl) acetate (900mg, 3.5mmol) in ethanol (10mL) was added a solution of sodium hydroxide (562mg, 14mmol) in water (10mL) and the reaction stirred at room temperature for 16h. The ethanol was then reduced in vacuo and the remaining aqueous solution was extracted with dichloromethane (2 x 30mL) and then acidified with 2M aqueous HCl (to pH 1). This solution was then extracted with ethyl acetate (2 x 30mL) and the combined organics were dried ($MgSO_4$) and reduced *in vacuo* to give 2-(4-ethylsulfonylphenyl) acetic acid (710mg, 88% yield). 1H NMR ($CDCl_3$): δH 1.28 (t, 3H), 3.12 (q, 2H), 3.77 (s, 2H), 7.51 (s, 2H) and 7.88 (s, 2H).

Step 3: To a solution of 2-(4-ethylsulfonylphenyl)acetic acid (700mg, 3.1mmol) in dichloromethane (20mL) was added N-(3-dimethylaminopropyl)-N-ethylcarbodiimide hydrochloride (705mg, 3.7mmol), 1-hydroxybenzotriazole hydrate (497mg, 3.7mmol) and 4-bromo-3,5-dichloroaniline (733mg, 3.1mmol) and the reaction was stirred at room temperature for 16h. The solution was then diluted with water (20mL) and extracted into dichloromethane (2 x 20mL). The organics were washed with 2M aqueous HCl solution (30mL), then saturated aqueous $NaHCO_3$ solution (30mL) and brine (30mL), dried ($MgSO_4$), reduced *in vacuo* and purified on the Biotage Isolera to give N-(4-bromo-3,5-dichloro-phenyl)-2-(4-ethylsulfonylphenyl)acetamide (520mg, 37% yield) as a light brown solid. 1H NMR (DMSO): δH 1.10 (t, 3H), 3.27 (q, 2H), 3.84 (s, 2H), 7.60 (d, 2H), 7.84-7.86 (m, 4H) and 10.64 (s, 1H). Step 4: A mixture of N-(4-bromo-3,5-dichloro-phenyl)-2-(4-ethylsulfonylphenyl)acetamide (100mg, 0.22mmol) , 2-(trifluoromethoxy)benzeneboronic acid (91mg, 0.4mmol), tetrakis(triphenylphosphine)palladium(0) (26mg, 0.02mmol) and potassium carbonate (61mg, 0.44mmol) in DMF (2mL) were reacted in the microwave at 100°C for 20min. After cooling to room temperature, the mixture was diluted with ethyl acetate (10mL) and washed with water (10mL). The organics were separated, dried, reduced *in vacuo* and purified on the Biotage Isolera to give N-[3,5-dichloro-4-[2-(trifluoromethoxy)phenyl]phenyl]-2-(4-ethylsulfonylphenyl)acetamide GSK2691805A (25mg, 21% yield) as an off-white solid. 1H NMR ($CDCl_3$): δH 1.32 (t, 3H), 3.15 (q, 2H), 3.84 (s, 2H), 7.24-7.27 (m, 1H), 7.36-7.42 (m, 3H), 7.46-7.50 (m, 1H), 7.55 (d, 2H), 7.66 (s, 2H) and 7.91 (d, 2H).
